# Spatiotemporal Modeling of GPCR Signaling: The Role of Endosomal Dynamics and Receptor Recycling

**DOI:** 10.64898/2026.04.29.721559

**Authors:** Chloé Weckel, Juliette Gourdon, Léo Darrigade, Vinesh Jugnarain, Pascale Crépieux, Eric Reiter, Frédéric Jean-Alphonse, Stefan Haar, Romain Yvinec

## Abstract

Cells communicate via extracellular ligands, such as hormones, which bind to plasma membrane receptors and trigger intracellular signaling cascades. G Protein-Coupled Receptors (GPCRs) exemplify this mechanism by initiating signaling both at the cell surface and, from intracellular compartments such as endosomes. The kinetics and spatial localization of these signals are critical determinants of cellular responses, yet receptor trafficking-including internalization, endosomal sorting, and recycling-remains a pivotal but often overlooked component of theoretical GPCR models. In this study, we present a mathematical framework that integrates receptor trafficking and signaling compartmentalization into generic GPCR dynamic models. Using a compartmentalized approach based on systems of ordinary differential equations (Chemical Reaction Networks), we analyze how receptor internalization and recycling modulate ligand-induced responses. Our results show that the balance between plasma membrane and endosomal signaling can significantly enhance or diminish ligand efficacy. Calibrated with high-throughput kinetic data, our model offers a refined tool for ligand pharmacological characterization and advances the understanding of GPCR signaling spatial organization.

## 1 Introduction

G protein-coupled receptors (GPCRs) constitute a large family of transmembrane receptors governing essential physiological functions, including reproduction, immune response, metabolism, and cardiovascular homeostasis. These receptors are targeted by a diverse array of ligands, such as hormones, ions, and neurotransmitters. Upon ligand binding, GPCR activation triggers intracellular signaling cascades that ultimately modulate gene expression and biological responses (Overington et al. 2006). Given their pivotal role in integrated physiological processes, GPCRs represent a major class of pharmacological targets. Classical pharmacological theory aims to quantify the action of both natural and synthetic ligands at a given receptor. The foundation of this field is the operational model, which draws an analogy between ligand-receptor-effector interactions and enzymatic processes (Black 1983). This model enables the quantification of ligand efficacy and potency for detecting ligand bias (Van Der Westhuizen et al. 2014; Stott et al. 2016), a promising concept for designing drugs with enhanced efficacy and reduced adverse effects. At steady state, the operational model establishes a relationship between ligand concentration and cellular response, facilitating dose-response analysis.

However, recent works have highlighted the critical influence of ligand binding kinetics (Klein Herenbrink et al. 2016) and receptor trafficking (Birtwistle and Kholodenko 2009) on cellular responses, giving rise to the concepts of kinetic and spatial bias in pharmacology (Hoare et al. 2020a; White et al. 2021). Indeed, the traditional view-where signaling cascades are triggered at the plasma membrane and subsequently desensitized via receptor internalization-has evolved substantially. For a growing number of GPCRs, it is now well established that receptors remain active in endosomes after their internalization and can trigger cellular responses distinct from those initiated at the plasma membrane (Calebiro et al. 2025). While the specific physiological role of this compartmentalized signaling has mostly been underappreciated, it has been elucidated for a few GPCRs. For the PTHR, cAMP produced at plasma membrane and at endosomes has been demonstrated to regulate distinct physiological functions (White et al. 2021). Similarly, signaling from endosomal compartments of the luteinizing hormone (LH) receptor has been shown to control the resumption of oocyte meiosis (preceding ovulation) (Lyga et al. 2016) and gonadal steroidogenesis (Gourdon et al. 2025).

The role of receptor trafficking-namely internalization into endosomes and recycling back to the plasma membrane-along with signaling from these compartments, calls for a revision of the standard operational model (Birtwistle and Kholodenko 2009). Receptor trafficking has been recognized as a key determinant of dose-response relationships. Deterministic frameworks based on ordinary differential equations (ODEs), derived from chemical reaction networks and structured through compartmentalized representations, have been developed to investigate these dynamics (Leelawattanachai et al. 2009; Birtwistle and Kholodenko 2009; Weddell and Imoukhuede 2017). Within this context, particular attention has been given to the characterization of unique steady states, which provide a compact description of cellular responses and support rigorous analysis of dose-response behaviors.

Since individual processes within GPCR signaling pathways are often experimentally inaccessible, estimating parameters for comprehensive models is a crucial step toward characterizing signaling compartmentalization. Importantly, it was recently reported that receptor intracellular distribution (e.g., among different endosomal subtypes) can be manipulated using innovative pharmacological tools (Raynaud et al. 2025). While previous studies have used trafficking models to estimate pharmacological quantities (efficacy, potency) and kinetic parameters (Birtwistle and Kholodenko 2009; Leelawattanachai et al. 2009), a systematic kinetic approach remains essential to dissect these mechanisms (Birtwistle and Kholodenko 2009; Hoare et al. 2018, 2020a,b).

The FSHR and the LHR play crucial roles in reproduction by supporting steroidogenesis and gametogenesis in both male and female. A distinctive feature of these gonadotropin receptors lies in their atypical endosomal distribution (Jean-Alphonse et al. 2014; Kalaidzidis et al. 2015; Sposini et al. 2017; Calebiro et al. 2025). Unlike many GPCRs that are internalized directly into Early Endosomes (EE), gonadotropin receptors are preferentially sorted into a distinct, smaller compartment known as Very Early Endosomes (VEE) (Jean-Alphonse et al. 2014; Sposini et al. 2017). This VEE pathway facilitates rapid receptor recycling to the plasma membrane and is established as a signaling-competent route. However, while recycling from VEEs is well-characterized, the fate of receptors located in EEs, and critically, their capacity to sustain signaling from this compartment, remain unclear. Hence, significant uncertainties persist regarding the precise mechanisms and kinetics governing the internalization, trafficking, and recycling of these receptors.

In this context, this study aims to elucidate how receptor trafficking (i.e., internalization and recycling) governs ligand-induced cellular responses. Using dose-response analyses, we show how trafficking dynamics and receptor spatial distribution shape cellular signaling outcomes. We further establish a framework for model selection and parameter estimation based on kinetic experiments conducted on the FSHR. Our results support the hypothesis that FSHR signals from two distinct endosomal compartments (e.g., VEEs and EEs), and we quantify the respective contribution of each compartment to the overall cellular response. Finally, we propose experimental strategies to reduce uncertainty in trafficking parameters.

## 2 Results

### 2.1 Receptor trafficking and spatial distribution shape cellular responses

The primary objective of this study was to quantify the impact of receptor internalization and recycling on second messenger production, given that it significantly influences cellular signaling. Throughout this paper, we focused on cyclic adenosine monophosphate (cAMP) as the second messenger, given its central role in the signaling cascades of *G*_*αs*_-coupled receptors and regulating central physiological functions. The framework presented here is broadly applicable to GPCRs and to signaling pathways mediated by distinct effectors. Given that blocking receptor internalization is the primary experimental method for studying the importance of post-endocytic signaling (Blythe et al. 2025), we specifically examined how internalization inhibition alters ligand-induced dose-responses. Our analysis revealed how such perturbations can uncover distinct trafficking and compartmentalization modes. Notably, this framework helped elucidate recent counterintuitive experimental findings across various GPCRs, where internalization inhibition was shown to affect ligand potency for some receptors while primarily impacting efficacy for others (Irannejad et al. 2015; Gourdon et al. 2025; Blythe et al. 2025).

Several assumptions underpin all models developed in this work and warrant brief introduction (see Methods 4.1, 4.2 for details). First, consistent with standard pharmacological modeling practices (Hoare et al. 2018, 2020a,b), we assumed that the ligand is in excess relative to the total receptor population during reversible binding reactions; consequently, the free ligand concentration was treated as constant. We developed models comprising either two or three signaling compartments based on ordinary differential equations (ODEs). The first compartment represents the plasma membrane, while the subsequent compartments could represent intracellular compartments such as Early Endosomes (EE), Very Early Endosomes (VEE), Golgi apparatus, mitochondria, or nucleus, depending on the specific GPCR (Suofu et al. 2017; Jiang et al. 2019; Nezhady and Rivera 2020; Crilly and Puthenveedu 2021). This framework was validated by considering FSHR as a model receptor, whose endosomal trafficking is atypical and remains poorly characterized to this day. In the context of FSHR, the intracellular compartment represents ‘endosomes’ in the two-compartments model. In the three-compartments model, these are explicitly distinguished as ‘very early endosomes’ (VEE) and ‘early endosomes’ (EE). These compartments were treated as fixed entities, with receptors capable of transitioning not only between the plasma membrane and intracellular spaces but also between distinct intracellular compartments, thereby establishing a fully compartmentalized system. Finally, we did not explicitly model ligand-receptor dissociation within intracellular compartments. Instead, we assumed that ligand dissociation occurs concomitantly with receptor recycling to the plasma membrane.

#### 2.1.1 Receptor trafficking critically modulates both the potency and efficacy of ligand-induced signaling

To investigate the effect of internalization and recycling on the production of cAMP, we first studied a model with two compartments, inspired from Birtwistle and Kholodenko (2009). Model 1 was developed to include reversible ligand-receptor binding, ligand-receptor internalization and receptor recycling, as well as catalytic cAMP production in both compartments and first-order cAMP degradation. Formulating model 1 as a system of ordinary differential equations (ODEs) (Methods 4.2.1-Eq. (12)), we described the temporal dynamics of the system and derived its unique steady-state for the cAMP response which reads, as a function of the total quantity of ligand (*L*), receptors (*R*_0_) and model parameters (Eq. (1)) (Birtwistle and Kholodenko 2009):

**Model 1.**
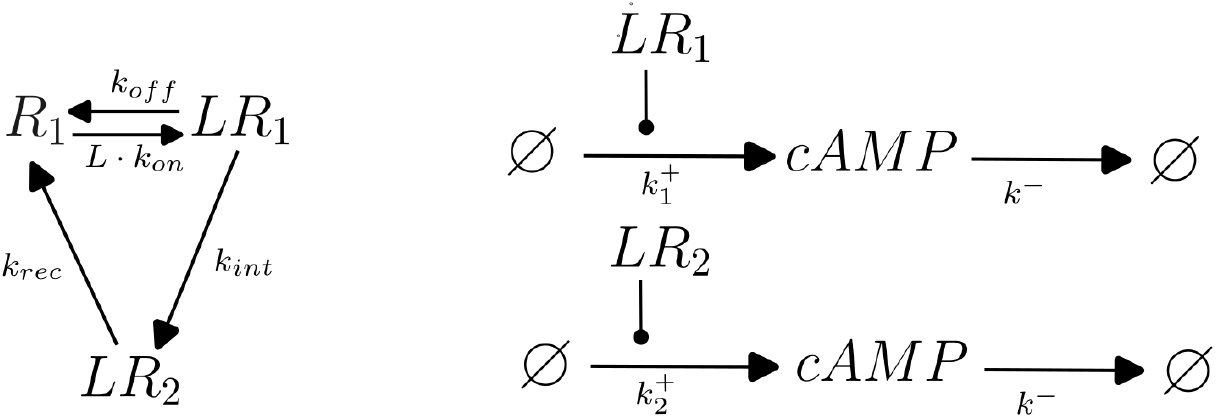
The left panel illustrates receptor trafficking, whereas the right panel depicts second messenger production and degradation. *R*_1_ denotes the amount of free receptors at the plasma membrane, *LR*_1_ and *LR*_2_ the amount of ligand-receptor complex at the plasma membrane and intracellular compartments. *cAMP* refers to the quantity of second messenger molecules.

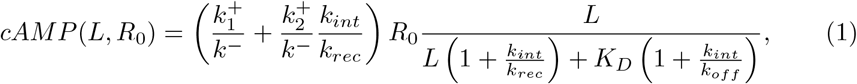

With 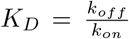. All calculations are detailed in Methods 4.2.1. From Eq. (1), we deduced that the efficacy is given by

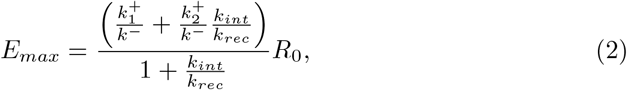

and the potency is determined by,

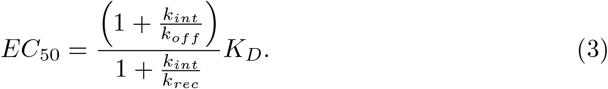

To investigate the effect of receptor trafficking, we compared expressions (1)–(3) with the classical ligand-receptor interaction model, when no internalization occurs (*k*_*int*_ = 0, Eq. (4)):

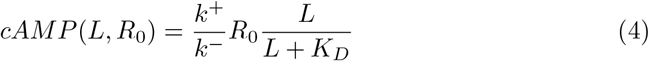

for which 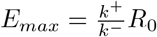 and *EC*_50_ = *K*_*D*_.

We thus deduced that receptor trafficking (*k*_*int*_, *k*_*rec*_) modulates both potency and efficacy of the signaling response, whereas second messenger production rates 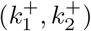 influence efficacy alone.

We subsequently focused our analysis on scenarios mimicking receptor internalization inhibition. As illustrated in Fig. 1, reducing the internalization rate can elicit diametrically opposite effects on the cellular response, depending on the relative magnitudes of signaling 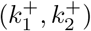 and receptor (*k*_*off*_, *k*_*rec*_) parameters. When the endocytic response is weaker than the plasma membrane response 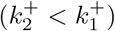, inhibiting receptor internalization increases efficacy (*E*_*max*_; Fig. 1a,b). Conversely, when the endocytic response is stronger than the plasma membrane response 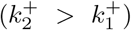, inhibiting receptor internalization decreases efficacy (*E*_*max*_; Fig. 1c,d).

**Figure 1.**
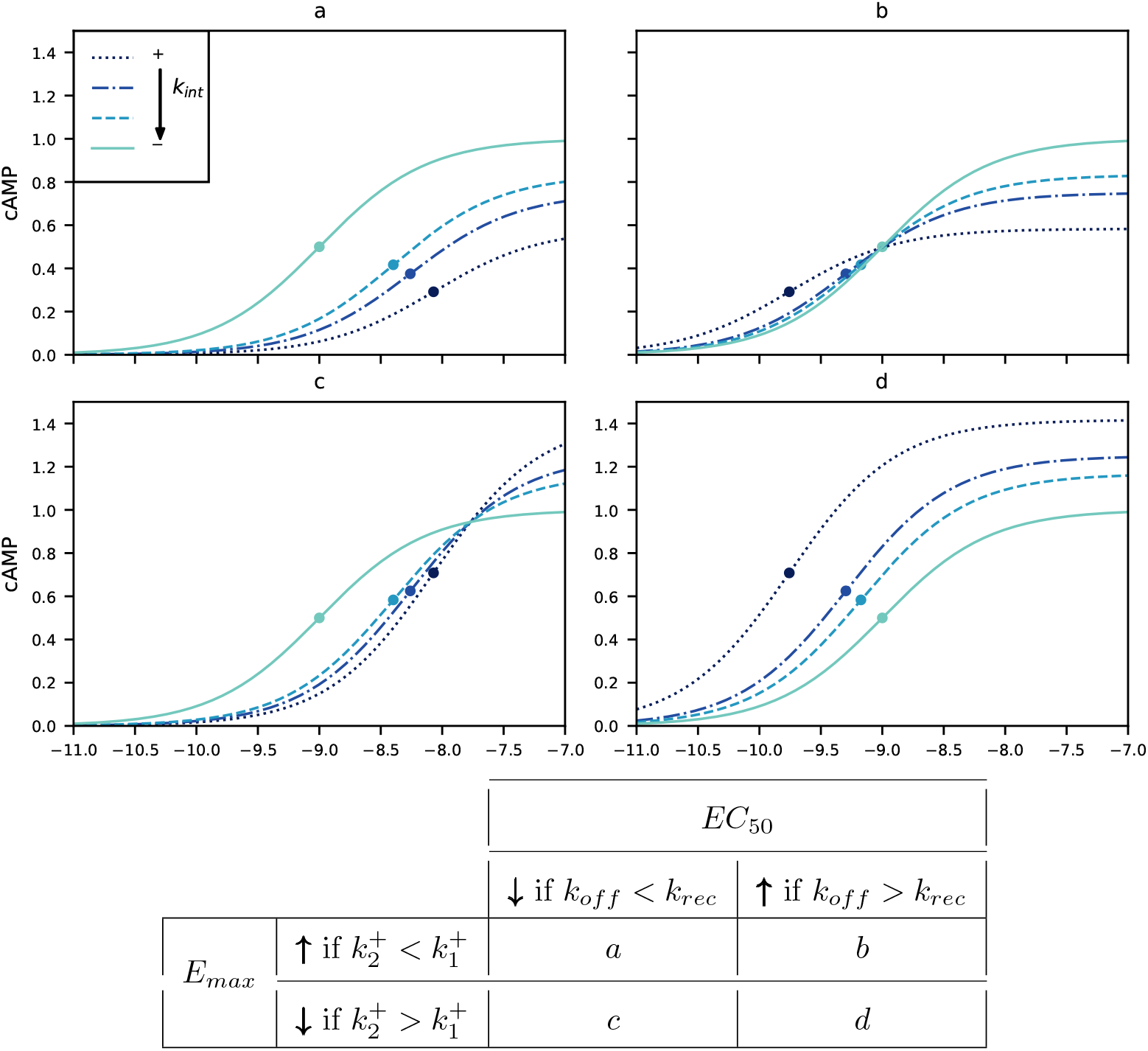
Impact of internalization inhibition on cAMP production using dose-response analysis. In the different panels, the FSH (M) dose response relationship of model 1 is plotted for different values of the internalization rate: *k*_*int*_ = 5 (dotted line), *k*_*int*_ = 1 (dash-dot line), *k*_*int*_ = 0.5 (dashed line), *k*_*int*_ = 0 (solid line). In each plot, circles represent the *EC*_50_ value. Common parameters to all six panels are *K*_*D*_ = 10^−9^, *R*_0_ = 1, *k*^−^ = 1, 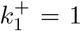, *k*_*rec*_ = 1. The remaining parameters values 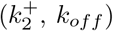 are given by: (a) 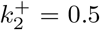, *k*_*off*_ = 0.1; (b) 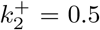, *k*_*off*_ = 100; (c) 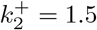, *k*_*off*_ = 0.1; (d) 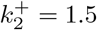, *k*_*off*_ = 100.

Similarly, when ligand dissociation from the receptor is faster than receptor recycling (*k*_*off*_ *> k*_*rec*_), inhibiting receptor internalization increases potency (*EC*_50_; Fig. 1b,d). In contrast, when ligand dissociation is slower than receptor recycling (*k*_*off*_ *< k*_*rec*_), inhibiting receptor internalization decreases potency (*EC*_50_; Fig. 1a,c). Reduction in ligand efficacy (Fig. 1c,d) is rather expected for receptors that have a strong intracellular activity, as inhibiting receptor internalization will prevent signaling from endosomal compartments. A decrease of ligand potency (Fig. 1b,d; increase of *EC*_50_) is more subtle, and shows that receptor trafficking may indeed affect the observed affinity from dose-response cellular assays. Going further, while inhibiting receptor internalization, the condition 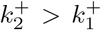 and *k*_*off*_ *> k*_*rec*_ (Fig. 1d) leads to a globally less efficient response (i.e. *E*_*max*_ decreases and *EC*_50_ increases), and symmetrically the condition 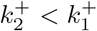 and *k*_*off*_ *< k*_*rec*_ (Fig. 1a) leads to a globally more efficient response (i.e. *E*_*max*_ increases and *EC*_50_ decreases). However, as shown in Fig. 1b,c, the effect of inhibiting receptor internalization may be more ambiguous. In these scenarios, efficacy and potency are affected in opposite directions. To resolve the complexities of cases (b) and (c) and characterize overall signaling efficiency under internalization blockade, we employed the transducer ratio *τ* (Eq. (5)), which defines coupling efficiency as the ratio of efficacy to potency:

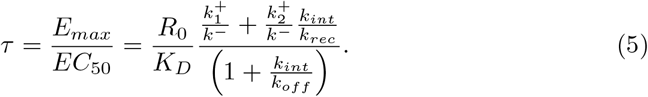

The transducer ratio (Eq. (5)) decreases when inhibiting receptor internalization if 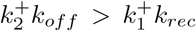 (Fig. 1d), and increases in the opposite case (Fig. 1a). However, the transducer ratio can increase or decrease in both cases (b) and (c). Given the widespread use of the transducer ratio to quantify ligand bias (Van Der Westhuizen et al. 2014), these results further suggest that it is also a reliable proxy for assessing how ligand-dependent responses are altered when internalization is perturbed.

Analytic formula (Eqs. (2),(3)) further reveal that the relative time scale of trafficking parameters (*k*_*int*_, *k*_*rec*_) and ligand unbinding rates (*k*_*off*_) affects potency exclusively, leaving efficacy unchanged. Indeed, if the trafficking parameter time scale decreases, with a constant ratio 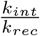, *E*_*max*_ remains unchanged, while the *EC*_50_ decreases. This indicates that, within this model, ligands inducing a slower receptor trafficking will have a tendency to have higher observed potency (Eq. (3)).

#### 2.1.2 Spatial distribution of receptors modulates both the potency and efficacy of ligand-induced signaling

The results derived in Eqs. (1)-(3) can be readily generalized to an arbitrary number of receptor-containing compartments. Beyond receptor trafficking kinetics, the spatial distribution of receptors within the cell may also influence the cellular response. To investigate this, we extended the analysis to model 2 with three compartments (e.g. very early and early endosomes in the case of gonadotropin receptors). Model 2 was constructed to account for reversible ligand-receptor binding, ligand-receptor internalization to two different compartments (*LR*_2_ and *LR*_3_) and receptor recycling from both intracellular compartments. It further incorporates compartment maturation, whereby ligand-receptor complexes *LR*_2_ transition to *LR*_3_. As in model 1, catalytic cAMP production occurs across all compartments, and cAMP degradation follows first-order kinetics. Interpreting again model 2 as an ODE system (Methods 4.2.2-Eq. (14)), we obtained a unique steady-state for the cAMP response, derived in Methods 4.2.2-Eq. (15), which yields analytical expressions for the pharmacological quantities *E*_*max*_ and *EC*_50_ (Methods 4.2.2-Eqs. (16)-(17)). In order, to investigate how the spatial distribution of ligand-receptor complexes influences the cellular response, we focused on conditions that promote their accumulation in the third compartment (Raynaud et al. 2025). To this end, we introduced a parameter *p*_23_ ∈ [0, 1], representing the fraction of internalized receptors directed to *LR*_3_ rather than *LR*_2_. The internalization rates were thus defined as *k*_*i*12_ = (1 − *p*_23_) · *k*_*int*_ and *k*_*i*13_ = *p*_23_ · *k*_*int*_ where *k*_*int*_ denotes the total internalization rate.

**Model 2.**
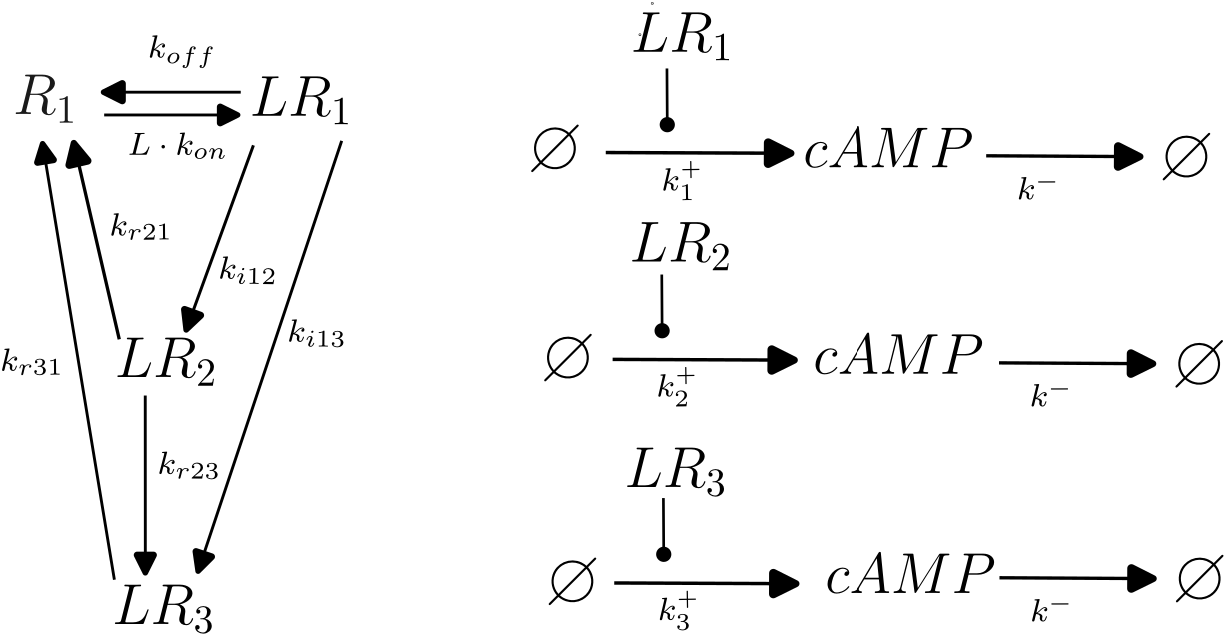
The left panel illustrates receptor trafficking, whereas the right panel depicts second messenger production and degradation. *R*_1_ denotes the amount of free receptors at the plasma membrane, *LR*_1_, *LR*_2_ and *LR*_3_ the amount of ligand-receptor complex at the plasma membrane and two types of intracellular compartments. *cAMP* refers to the quantity of second messenger molecules.

From the analytical formula for the steady-state cAMP response (Methods 4.2.2-Eq. (15)), we obtained the efficacy, potency and transducer ratio values as follows:

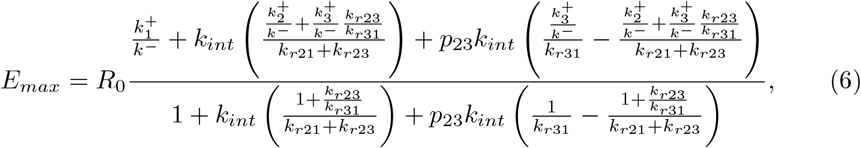

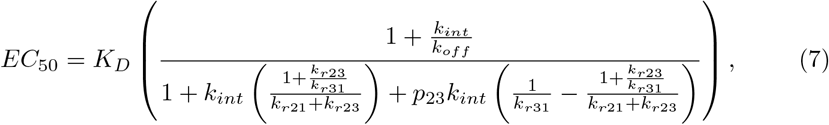

and

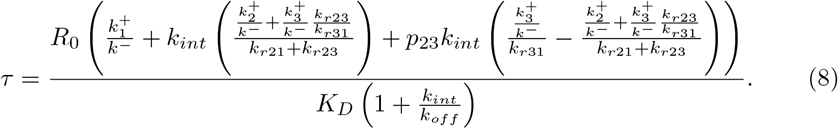

As in Fig. 1, Fig. 2 illustrates that promoting ligand-receptor accumulation in the third compartment (*e*.*g*. increasing *p*_23_) can exert opposing effects on potency, efficacy, and transducer ratio. This reflects the intricate interplay between receptor trafficking parameters and signaling parameters, as captured by Eqs. (6)-(8). Remarkably, inducing ligand-receptor accumulation in the third compartment (e.g. increasing *p*_23_) always leads to a monotonic relation between *E*_*max*_, *EC*_50_ and *τ* with respect to *p*_23_. However, as shown on Fig. 2, all qualitatively distinct cases are theoretically attainable. Characterizing the dynamics of each case could help identify the most plausible scenario based on current knowledge, and provide a framework for dissecting receptor trafficking and compartmentalized signaling. Notably, the compartment maturation rate from *LR*_2_ to *LR*_3_ (*k*_*r*23_) has no effect on the variations of the potency, efficacy or transduction coefficient. Potency is affected only by recycling rates, consistent with the result obtained for the first model. The interpretation of the first case (Fig. 2a) is as follows: when the recycling rate is faster from *LR*_2_ than from *LR*_3_ (*k*_*r*21_ *> k*_*r*31_), and cAMP production in the third compartment is high enough (Condition 18), then the accumulation of receptor in the third compartment leads to globally more efficient response (*E*_*max*_ increases, *EC*_50_ decreases, *τ* increases). The fourth case (Fig. 2d) is symmetrically opposite to the first case, with a globally less efficient response.

**Figure 2.**
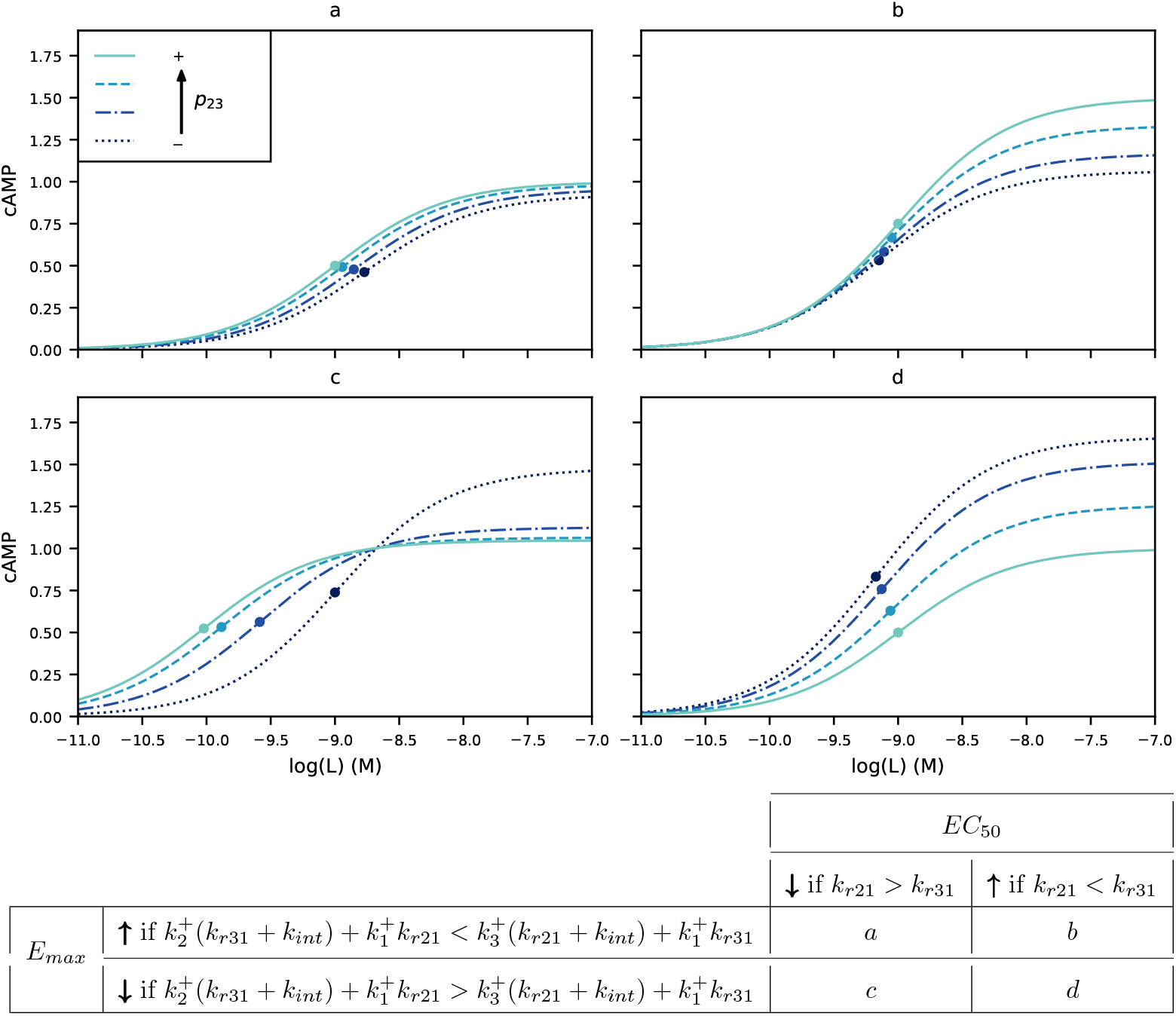
Impact of biasing receptor distribution on cAMP production using dose-response analysis. Circles represent the value of *EC*_50_. The values of *p*_23_ represent the increasing of receptors in the third compartment. Common parameters are *p*_23_ = 0 (dotted line), *p*_23_ = 0.3 (dashdot line), *p*_23_ = 0.7 (dashed line) and *p*_23_ = 1 (solid line), ; *K*_*D*_ = 10^−9^, *R*_0_ = 1, *k*^−^ = 1, 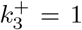, *k*_*int*_ = 1, *k*_*r*23_ = 1, *k*_*off*_ = 1, whereas the recycling (*k*_*r*21_ and *k*_*r*31_) and production (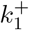 and 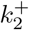) parameters are specific to each panel: (a) *k*_*r*21_ = 10, *k*_*r*31_ = 1, 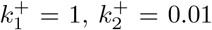; (b) *k*_*r*21_ = 0.01, *k*_*r*31_ = 1, 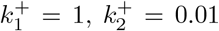; (c) *k*_*r*21_ = 20, *k*_*r*31_ = 0.05, 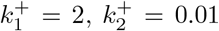; (d) *k*_*r*21_ = 0.01, *k*_*r*31_ = 10, 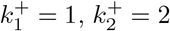.

In contrast, the second (Fig. 2b) and third cases (Fig. 2c) are more subtle. In those later cases, the impact of receptor accumulation in the third compartment is dose-dependent, and the overall effect can be quantified by the transduction coefficient *τ*, whose variation depends on the relation between signaling efficacy and receptor recycling (Condition 19).

In conclusion, unlike the receptor internalization inhibition case studied in Subsect. 2.1.1, a change in *E*_*max*_ induced by biasing receptor spatial distribution is harder to interpret in terms of parameters values and, therefore, receptor trafficking or cAMP production. However, a shift in the dose-response curve (*EC*_50_) is indicative of differential recycling rates from distinct intracellular compartments, and the transduction coefficient *τ* may further provide insight into the signaling efficacy within each compartment.

### 2.2 Evidence that FSHR signals through distinct compartments

As demonstrated by the results presented in the previous section, receptor trafficking and localized signaling parameters substantially influence cellular responses. The primary objective was therefore to identify the parameter regime governing a given signaling pathway from biological data. To this end, we developed a workflow spanning from model design to prediction, encompassing several intermediate steps. This workflow was inspired by (Villaverde et al. 2022; Banga and Villaverde 2025) and leveraged StructuralIdentifiability (Dong et al. 2023), PEtab (Schmiester et al. 2021), and pyPESTO (Schälte et al. 2023) to perform parameter estimation from kinetic experiments developed on FSHR (Methods 4.4). FSHR was considered as a model receptor, whose endosomal trafficking is atypical. Receptors are first internalized into a compartment called the very early endosome (VEE), and then into a compartment called the early endosome (EE). However, the spatial organization of FSHR remains poorly characterized to this day.

#### 2.2.1 Dose-response kinetic experiments and model design

Using Bioluminescence Resonance Energy Transfer (BRET) measurements, we monitored the temporal dynamics of molecular species following FSH stimulation in a cell system overexpressing FSHR (Fig. EV1) (Pearce et al. 2025). We considered four different BRET experiments. First, we quantified total cAMP levels using an intramolecular biosensor (Methods 4.11.2) upon continuous stimulation with varying FSH concentrations. At one FSH concentration (1 nM), cAMP level was monitored in control conditions or in the presence of PitStop2, a chemical inhibitor of clathrin-mediated endocytosis (CME), hereafter referred to as *ps*_2_. The variable representing cAMP quantity is denoted *cAMP*, and 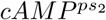 when the internalization inhibitor is present. Localization of receptor in specific cellular compartments was assessed by monitoring interaction between FSHR fused to the luciferase RLuc8 (FSHR-RLuc8) and a fluorescent protein Ypet localized either at plasma membrane (LYN-Ypet) or at early endosomes (Ypet-FYVE) (Fig. EV2). Measuring the interaction between LYN-Ypet and FSHR-RLuc8 enables to monitor FSHR localization at the plasma membrane, which decrease upon ligand-induced internalization. In contrast, monitoring interaction between FSHR-RLuc8 and Ypet-FYVE allowed to quantify the time-dependent accumulation of ligand-receptors complexes in early endosomes. Drawing on prior knowledge of gonadotropin receptors trafficking (Jean-Alphonse et al. 2014; Sposini et al. 2017) and accounting for remaining uncertainties regarding receptor trafficking mechanisms, we first designed a complete “maximal” model (Model 3), analogous to Model 2, in which *LR*_1_, *LR*_2_ and *LR*_3_ represent ligand-receptor complexes at the plasma membrane (PM), in the very early endosomes (VEE) or early endosomes (EE) (Fig. EV2). We further incorporated the possibility of irreversible receptor desensitization from *LR*_1_, at the plasma membrane, and from *LR*_3_ at the early endosomes (Model 3). This complete model yielded an ODE system detailed in Methods 4.2.3-Eq. (20).

We assumed the BRET signal is proportional to the quantity of species in the cell (Tripp et al. 2022). The BRET signals then relates to cAMP levels, plasma membrane receptors (*PM*_*R*_), and ligand-receptor complexes within EE (*EE*_*R*_).

**Model 3.**
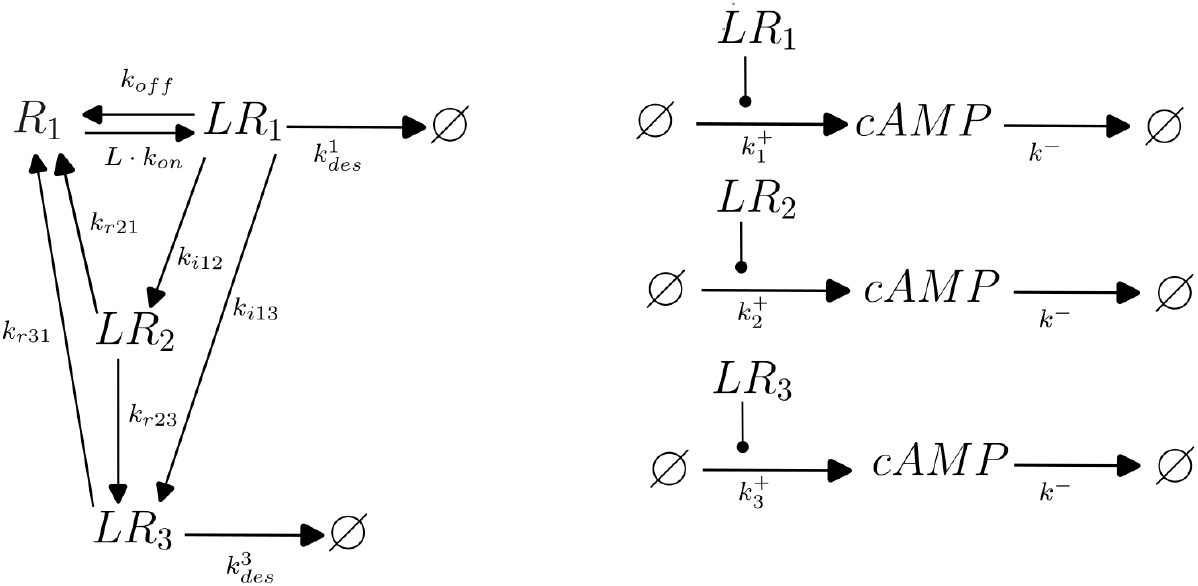
The left panel illustrates receptor trafficking, whereas the right panel depicts second messenger production and degradation. *R*_1_ denotes the amount of free receptors at the plasma membrane, *LR*_1_, *LR*_2_ and *LR*_3_ the amount of ligand-receptor complex at the plasma membrane (PM) and two types of intracellular compartments. *cAMP* refers to the quantity of second messenger molecules.

These data were linked to species evolution according to the following observable equations:

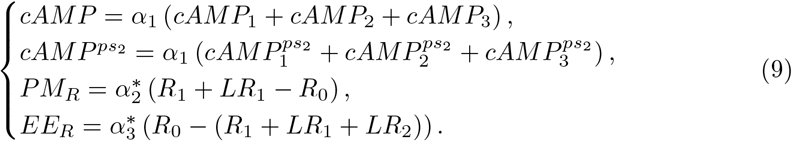

Kinetic cAMP measurements were performed at varying ligand concentrations (*FSH* = 0, 0.1, 0.3, 1, 3, 10 nM). Measurements in the presence of the clathrin-mediated endocytosis inhibitor PitStop2 were performed at a single FSH concentration (*FSH* = 1 nM). Finally, receptor internalization (*PM*_*R*_) and trafficking to the EEs (*EE*_*R*_) kinetics were assessed in basal conditions (*FSH* = 0) and with *FSH* = 100 nM.

Importantly, prior to parameter estimation, it was essential to verify that the experimental design and model structure yielded a well-posed - i.e., identifiable - inverse problem. To this end, we first reparametrized the ODE system, together with the observable equations, and non-dimensionalize the parameters, so as to reduce the total number of parameters as much as possible (Methods 4.3). The resulting normalized model leads to Eq. (21), comprising 19 parameters in which new parameters are called *λ*_*tot*_, *λ*_1_ and *λ*_3_ and representing the cellular response produced by receptors, Eqs. (22))(Table 3) instead of 21 (Table 2). Applying the methodology of (Dong et al. 2023), we demonstrated that the parameter estimation problem is locally identifiable given the observables (Eq. (9)) meaning that each set of observable outputs locally determines a single set of parameter values (Methods 4.6). Interestingly, when the internalization inhibition data with PitStop2 were excluded, the model became non identifiable for a subset of parameters, including several trafficking and cellular response parameters (*λ*_*tot*_, *k*_*lr*13_, *k*_*r*21_, *k*_*r*31_, *k*_*r*21_, 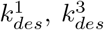, *λ*_1_ and *λ*_3_). This result demonstrates the value of the experimental design in jointly constraining - and thereby disentangling - receptor trafficking and compartmentalized signaling parameters.

**Table 1.** Averaged outputs for the 13 best models. The table presents the proportion of internalized receptors (*LR*_2_ + *LR*_3_) relative to total receptors, and the proportion of receptors in VEE or EE relative to internalized receptors (*LR*_2_ + *LR*_3_), but also endocytosis and recycling time and probability for a 5min incubation with 10nM of ligand (Methods 4.10). Values in brackets indicate uncertainty.

| Model 3. | 1 | 2 | 3 | 4 |  |
| --- | --- | --- | --- | --- | --- |
| $LR_{intra}$ (%) | 0.05<br>[0.04, 1.15] | 0.05<br>[0.04, 1.71] | 0.01<br>[0.00, 1.27] | 0.01<br>[0.00, 1.21] | |
| $LR_2$ (%) | 9.98<br>[2.16, 25.34] | 96.75<br>[87.36, 97.75] | 68.55<br>[2.34, 98.85] | 85.22<br>[2.17, 98.90] | |
| $LR_3$ (%) | 90.02<br>[74.66, 97.84] | 3.25<br>[2.25, 12.64] | 31.45<br>[1.15, 97.66] | 14.78<br>[1.10, 97.83] | |
| Time endocytosis (s) | 55.97<br>[48.66, 65.21] | 55.83<br>[51.02, 60.20] | 55.67<br>[9.41, 65.51] | 55.64<br>[9.56, 65.18] |  |
| Probability endocytosis | 0.01<br>[0.00, 0.14] | 0.01<br>[0.01, 0.24] | 0.01<br>[0.00, 0.84] | 0.01<br>[0.01, 0.83] |  |
| Time recycling (s) | 78.94<br>[47.91, 227.85] | 61.42<br>[32.19, 77.47] | 4.43<br>[0.03, 95.78] | 0.45<br>[0.02, 0.85] |  |
| Probability recycling | 1.00<br>[1.00, 1.00] | 1.00<br>[1.00, 1.00] | 1.00<br>[1.00, 1.00] | 0.98<br>[0.00, 1.00] |  |
| Model 3. | 5 | 6 | 7 | 8 |  |
| $LR_{intra}$ (%) | 0.01<br>[0.01, 1.25] | 0.01<br>[0.01, 1.29] | 0.05<br>[0.04, 1.25] | 0.05<br>[0.04, 1.17] | |
| $LR_2$ (%) | 49.74<br>[0.36, 99.76] | 51.81<br>[0.36, 99.77] | 9.99<br>[2.17, 25.25] | 9.99<br>[2.17, 26.79] | |
| $LR_3$ (%) | 50.26<br>[0.24, 99.64] | 48.19<br>[0.23, 99.64] | 90.01<br>[74.75, 97.83] | 90.01<br>[73.21, 97.83] | |
| Time endocytosis (s) | 55.63<br>[9.55, 64.39] | 55.63<br>[9.68, 63.56] | 55.97<br>[49.30, 63.93] | 55.98<br>[49.32, 65.62] |  |
| Probability endocytosis | 0.01<br>[0.01, 0.83] | 0.01<br>[0.01, 0.83] | 0.01<br>[0.00, 0.15] | 0.01<br>[0.00, 0.14] |  |
| Time recycling (s) | 7.64<br>[0.03, 76.63] | 7.12<br>[0.03, 84.66] | 78.92<br>[47.33, 95.60] | 78.95<br>[47.84, 229.13] |  |
| Probability recycling | 1.00<br>[1.00, 1.00] | 0.92<br>[0.04, 1.00] | 1.00<br>[0.00, 1.00] | 1.00<br>[1.00, 1.00] |  |
| Model 3. | 9 | 10 | 11 | 12 | 13 |
| $LR_{intra}$ (%) | 0.05<br>[0.04, 1.22] | 0.05<br>[0.04, 1.71] | 0.05<br>[0.04, 1.72] | 0.05<br>[0.04, 1.72] | 0.05<br>[0.04, 1.71] |
| $LR_2$ (%) | 9.08<br>[0.33, 23.94] | 96.75<br>[87.36, 97.80] | 96.75<br>[87.36, 98.84] | 96.75<br>[87.07, 98.47] | 96.75<br>[87.36, 98.71] |
| $LR_3$ (%) | 90.92<br>[76.06, 99.67] | 3.25<br>[2.20, 12.64] | 3.25<br>[1.16, 12.64] | 3.25<br>[1.53, 12.93] | 3.25<br>[1.29, 12.64] |
| Time endocytosis (s) | 55.97<br>[48.97, 64.84] | 55.82<br>[50.21, 60.92] | 55.83<br>[49.36, 62.82] | 55.83<br>[49.44, 65.40] | 55.83<br>[51.02, 65.30] |
| Probability endocytosis | 0.01<br>[0.01, 0.15] | 0.01<br>[0.01, 0.24] | 0.01<br>[0.01, 0.24] | 0.01<br>[0.01, 0.24] | 0.01<br>[0.01, 0.24] |
| Time recycling (s) | 78.86<br>[47.76, 245.29] | 61.49<br>[34.29, 72.82] | 61.42<br>[34.26, 74.47] | 61.42<br>[20.43, 74.92] | 61.42<br>[32.19, 73.46] |
| Probability recycling | 1.00<br>[1.00, 1.00] | 1.00<br>[1.00, 1.00] | 1.00<br>[1.00, 1.00] | 1.00<br>[1.00, 1.00] | 1.00<br>[1.00, 1.00] |

**Table 2.** Table of the parameters of model 3.

| Type | Name | Number |
| --- | --- | --- |
| Initial conditions | $R_0$ | 1 |
| Kinetic parameters | $k_{on}, k_{off}, k_{r21}, k_{r31}, k_{i12}, k_{i13}, k_{r23}, k_1^+, k_2^+, k_3^+, k^-, k_{des}^1, k_{des}^3$ | 13 |
| Drug parameter | $ps_2$ | 1 |
| Error parameters | $\sigma_1, \sigma_2, \sigma_3$ | 3 |
| Data parameters | $\alpha_1, \alpha_2, \alpha_3$ | 3 |
| Total |  | 21 |

**Table 3.** Table of the reparameterized parameters (Model 3 and Eq. (21)).

| Type | Name | Number |
| --- | --- | --- |
| Time scale parameter | $k_{off}$ | 1 |
| Kinetic parameters | $K_D, k_{r21}^*, k_{r31}^*, k_{i12}^*, k_{i13}^*, k_{r23}^*, \lambda_{tot}, \lambda_1, \lambda_3, k^{*-}, k_{des}^{1*}, k_{des}^{3*}$ | 12 |
| Drug parameter | $ps_2$ | 1 |
| Error parameters | $\sigma_1, \sigma_2, \sigma_3$ | 3 |
| Data parameters | $\alpha_2, \alpha_3$ | 2 |
| Total |  | 19 |

#### 2.2.2 Identifying a cluster of models

We developed a model selection procedure aimed at identifying parsimonious models with fewer parameters than the complete model 3, in order to discriminate among the following alternative questions, given the available data (Fig. 3):

**Figure 3.**
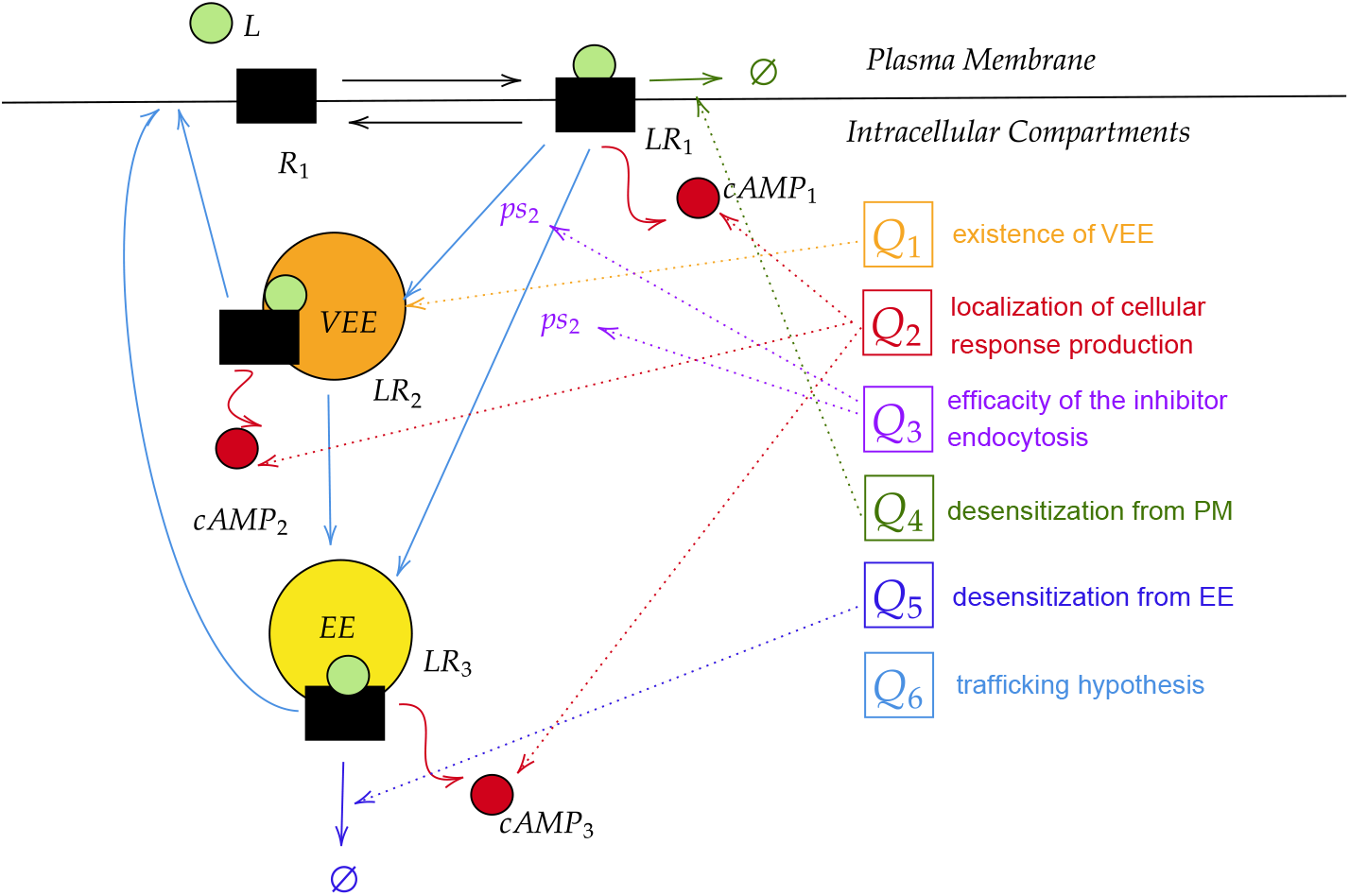
The different questions for the model selection. Questions refers to Q1-Q6. Q1. Does the ligand-receptor complex internalize in one or two compartments (EE or VEE/EE) i.e., is *k*_*i*12_ = 0? Q2. Is the ligand-receptor complex functionally active in two compartments (PM and VEE, i.e., is 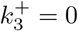 or PM and EE, i.e., is 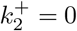) or three compartments (PM, VEE and EE)? Q3. Is chemical internalization inhibition partially or fully efficient, i.e. is *ps*_2_ = 0? Q4. Are there irreversible receptor desensitization mechanisms at the plasma membrane, i.e., is 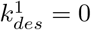 Q5. Are there irreversible receptor desensitization mechanisms at early endosomes, i.e., is 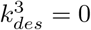 Q6. If there are three active signaling compartments, what is the trafficking pathway that the ligand-receptor complex may follow (internalization, traffic to an other compartment, recycling)– i.e. what is the most plausible trafficking model among 8 competitive models (Fig. EV3)?

Those six questions lead to 200 submodels of the complete Model 3. We ran the model selection algorithm (Methods 4.5) on the 200 models (Appendix Table S2). Rather than a single best model, a cluster of models was identified according to the Δ_*AIC*_ criterion (Burnham and Anderson). 13 models have a Δ_*AIC*_ *<* 2 and are therefore considered as likely as be the best model; moreover, up to 35 models have 2 *<* Δ_*AIC*_ *<* 7, and can thus be regarded as suitable alternatives. Importantly, all thirteen best models share common features. First, three compartments are required (Q1), with ligand-receptor complexes being functionally active either within each compartment individually or within two compartments: PM and EE (*LR*_1_ and *LR*_3_) (Q2). Model selection rejects the scenario in which production arises solely from PM and VEE (*LR*_1_ and *LR*_2_), even though biological model suggests that active receptors may be more abundant in VEE (*LR*_2_) than in EE (*LR*_3_). The data are compatible with the hypothesis that the CME inhibitor is fully effective (Q3) ^1^ and with the existence of an irreversible receptor desensitization mechanism at the plasma membrane (Q4). An irreversible desensitization mechanism from early endosomes is required only when recycling from this third compartment is absent, presumably to prevent intracellular receptor accumulation (Q5). Finally, we cannot currently identify a unique best trafficking model (Q6), as the eight tested (Fig. EV3) fall below the threshold Δ_*AIC*_ *<* 2 (Appendix Table S2). We therefore verified that the 13 best models yield comparable fits, as illustrated in Fig. 4, which demonstrates strong agreement between model predictions and experimental data across all observables - cAMP production, receptor internalization, and receptor trafficking to early endosomes - both under control conditions and upon CME inhibition.

**Figure 4.**
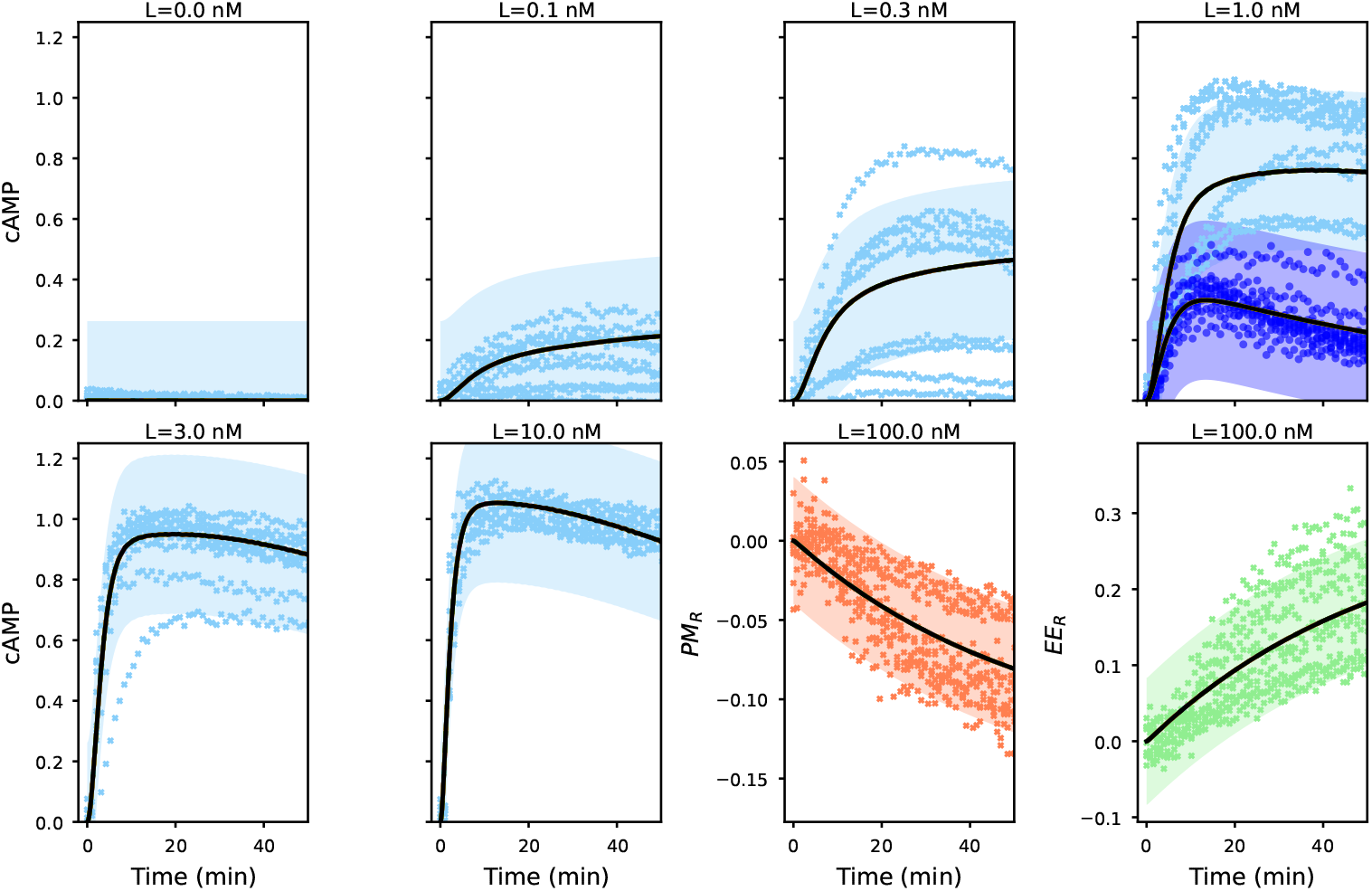
Data fitting with the thirteen best models. The thirteen best fit of the ODE simulation (Appendix Table S2, models 1-13) are superimposed while shading indicating statistical uncertainty for the first model (Methods 4.7). Experimental data points are shown as crosses. For the 1 nM ligand condition, the bottom curve (shown in dark blue) corresponds to cAMP levels measured under PitStop2 treatment.

Verifying practical parameter identifiability for the different models is essential to perform reliable model prediction. Hence, to further investigate the thirteen best models (Appendix Table S2, models 1-13), we performed a profile likelihood estimate for each parameter on each model (Methods 4.6).

Some parameters are clearly identifiable with very small confidence intervals and common values across the eight models (*k*_*off*_, *K*_*D*_, *k*^−∗^, *σ*_1_, *σ*_2_, *σ*_3_, 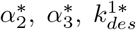), whereas some are consistent across models but have larger confidence intervals (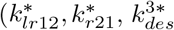, *λ*_1_, *λ*_*tot*_). The remaining parameters are consistent across some models but exhibit large confidence intervals in others (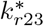, *λ*_3_, *k*_*i*13_, 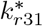) (Fig. 5). Parameters governing time scales, ligand-receptor interactions, and observables are well identified, while those related to receptor trafficking and localized cAMP production remain subject to greater uncertainty. In addition, two consistent groups emerge when considering the models: those in which receptors remain active in all compartments (models 3.1, 3.3, 3.4, 3.5, 3.6, 3.7, 3.8, 3.9), and those in which receptor activity is restricted to the plasma membrane and early endosomes (*LR*_1_ and *LR*_3_) (models 3.2, 3.10, 3.11, 3.12, 3.13). This distinction is particularly evident for several parameters, including 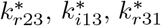, *λ*_3_ (Fig. 5). These two groups will be compared in the subsequent study, in particular by considering models 3.1 and 3.2, the two first selected models.

**Figure 5.**
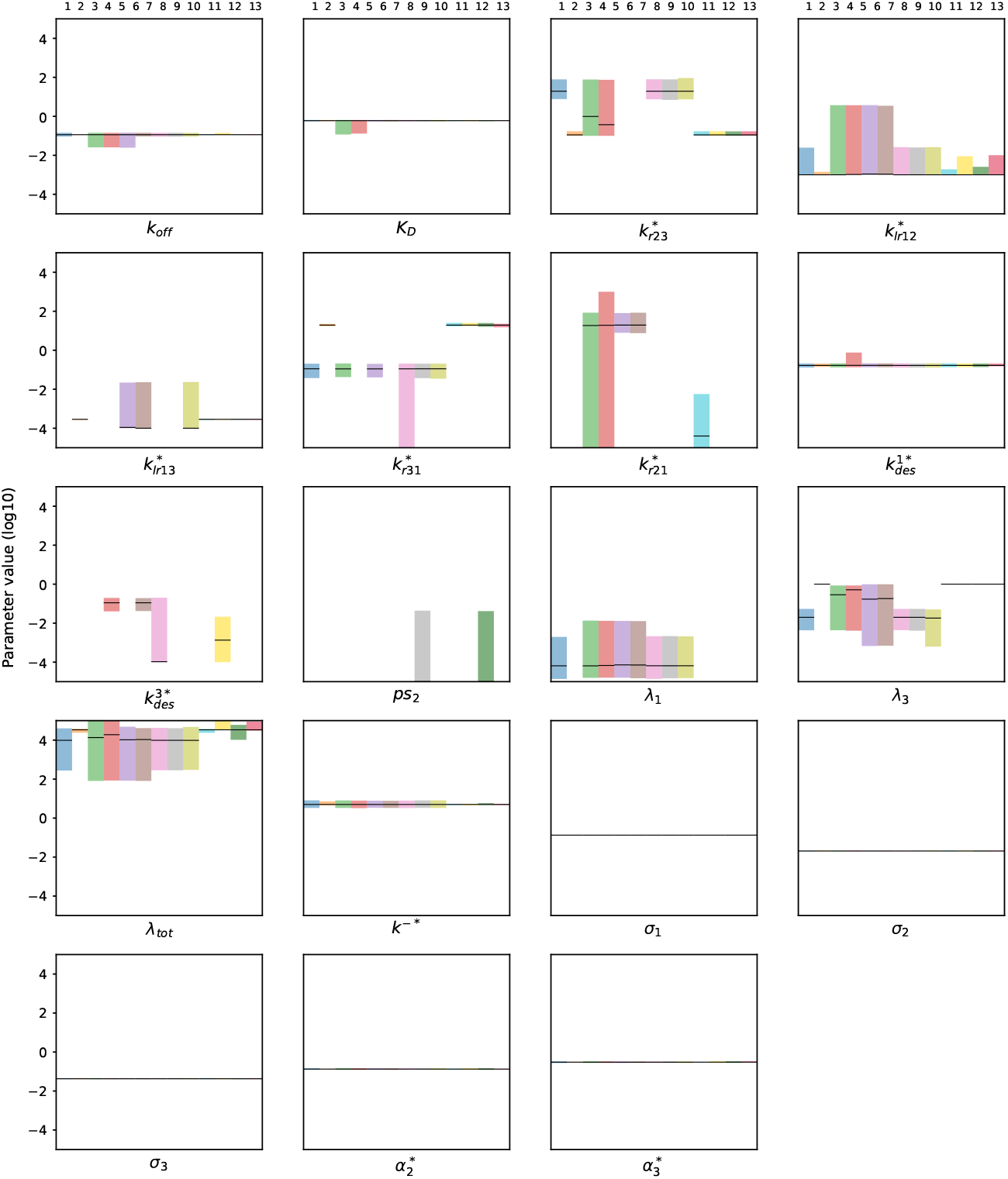
Confidence intervals for all parameters in the 13 first models. In each panel, we show the *profile-based* 95% confidence interval (Methods 4.7) of a given parameter for the 13 selected best models (Appendix Table S2). The first line corresponds to the selected model, 3.x, where x denotes the number of models from 1 to 13. The optimum value is represented by the dark line. arameters noticed with *k*^∗^ are the normalized parameters: 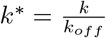 and 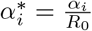.

### 2.3 Model prediction and uncertainty quantification

#### 2.3.1 The remaining uncertainties within the best thirteen models concern the amounts of receptor complexes in the intracellular compartments

Of note, parameter uncertainty may arise from the inherent variability of the experimental data and/or from an insufficient number of observables or experimental conditions. As apparent from Fig. 4, experimental variability is high, which necessarily increases the variance of the statistical error model. The 13 selected models fall into two groups: those with cAMP production from ligand-receptor complexes in all three compartments (Fig. 6-first two columns), and those with production restricted to two compartments, *LR*_1_ and *LR*_3_ (Fig. 6-third and fourth columns). Parameter uncertainty with respect to the cellular response is greater in the latter case. In Fig. 6 (first and third columns), we plot the *profile-based* 95% confidence interval (shaded area), revealing that parameter uncertainty does not affect cAMP level predictions for model 3.1, in contrast to model 3.2, but significantly impacts the predicted amount of internalized receptors. Similarly, Fig. 6 (second and fourth columns) shows that model predictions for each cAMP variable are consistent across the best models in both groups. However, whereas models within the first group differ substantially in their predictions of internalized receptor amounts, these quantities remain consistent across all models in the second group, where cAMP are produced solely from receptors active at the PM and EE (*LR*_1_ and *LR*_3_).

**Figure 6.**
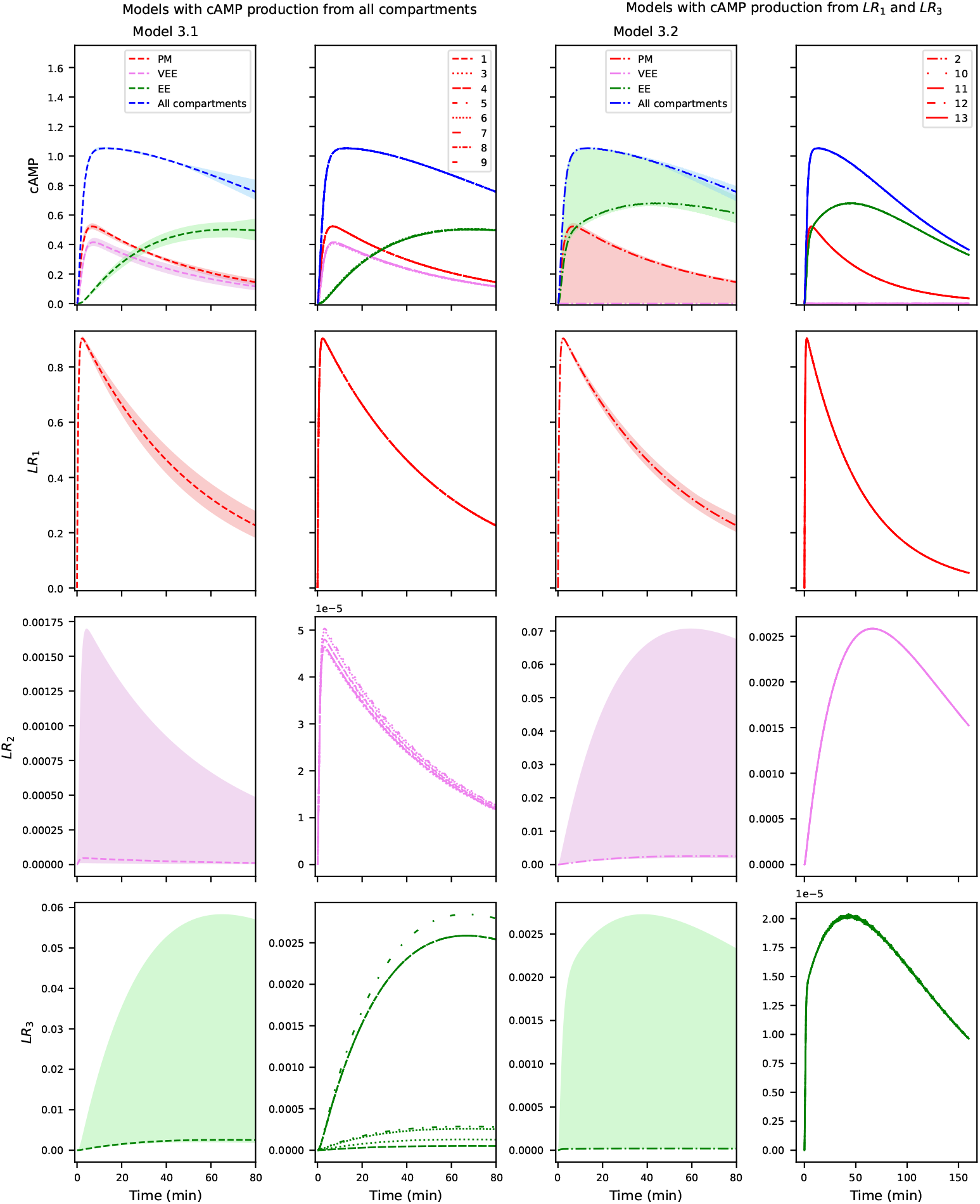
Prediction and uncertainty quantification of cAMP levels and LR complexes in each compartment for the 13 best models. Model 3.1 and model 3.2 predictions are shown with 95% profile-based confidence intervals (Methods 4.7), alongside single predictions for all best models with ligand-receptor complex production from all compartments (columns 1 and 2) or from *LR*_1_ and *LR*_3_ (columns 3 and 4) in response to a single FSH dose (10 nM). Parameters correspond to the best estimates obtained using the AIC criterion (Appendix Table S3).

Despite our theoretical identifiability results, the observed prediction uncertainty in internalized receptor amounts likely reflects uncertainty in trafficking and localized production parameters, which may partially compensate for one another to reproduce a given cAMP output-at least within our dataset. A negative correlation can indeed be observed between the number of ligand-receptor complexes in EE (*LR*_3_) and the cAMP production rate in this compartment 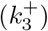. The amount of *LR*_3_ is higher for models with trafficking hypotheses *c* and *e*, while cAMP production in the third compartment is lower in these models (Fig. 6, Appendix Table S3). These two models have no recycling from the second compartment, *LR*_2_, meaning that recycling occurs only from the third compartment, *LR*_3_, which ultimately increases the total number of *LR*_3_ complexes. The proportion of internalized receptors distributed across intracellular compartments varies between the 13 models (Table 1): from 10% to 97% for *LR*_2_, and from 3% to 90% for *LR*_3_. Consistently, the highest *LR*_2_ values are observed in models in which no cAMP is produced from receptors in the second compartment, VEE-*LR*_2_ (models 3.2, 3.10, 3.11, 3.12, and 3.13). In contrast, when cAMP is produced from receptors in both compartments, *LR*_3_ exceeds *LR*_2_ (models 3.1, 3.7, 3.8, and 3.9), despite the same proportion of intracellular compartments in these two groups (0.05%). Uncertainties are larger for the four models with a balanced receptor distribution between compartments and a lower proportion of intracellular compartments (0.01%) than in the first two groups (models 3.3, 3.4, 3.5, and 3.6). To further discriminate the trafficking models among the best models, an interesting perspective would be to design experiments to monitor the quantities of *LR*_2_ and *LR*_3_ using specific sensors that are selectively localized in these compartments (Jean-Alphonse et al. 2014).

Furthermore, designing localized sensors to monitor either receptors or second messenger molecules in a specific compartment would strengthen our conclusions regarding hypotheses Q1 to Q5, as such data would enable further discrimination between models. To illustrate this, Fig. EV4 shows model predictions for models 3.43, 3.48, 3.68, and 3.124, which all share the common *c* trafficking structure as their best model (Fig. EV3), but differ in their assumptions regarding hypotheses Q5, Q4, and Q2, respectively (Appendix Table S2). These alternative models deviate only marginally from the best model in terms of total cAMP - failing in particular to capture the long-time cAMP decrease - but diverge more substantially in their predictions of internalized receptor amounts and endosomal cAMP levels (Appendix Table S4).

#### 2.3.2 FSHR activation leads to a strong endosomal signaling response

All the best models exhibit comparable behavior, with very few internalized receptors (less than 0.05%, Table 1) that elicit a strong cellular response (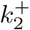 and 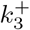 are three to four order of magnitude higher than 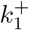, Appendix Table S3). The first and transient wave of cellular response originates from signaling at the plasma membrane and in very early endosomes (*LR*_2_), with a slightly higher response in VEE (models 3.1, 3.3, 3.4, 3.5, 3.6, 3.7, 3.8, 3.9). A second and sustained wave of signaling corresponds to cAMP production in EEs (*LR*_3_) (Fig. 6). Further, receptor internalization rates (of order 10^−4^ − 10^−5^*s*^−1^) appear to be considerably slower than receptor recycling rates (of order 10^−2^ − 10^0^*s*^−1^), the latter being comparable to ligand unbinding rate (of order 10^−1^*s*^−1^). All models predict an averaged endocytosis time of 56*s* with a rather sharp confidence interval, while the averaged receptor recycling time has much more uncertainty across models, ranging from 0.5 to 79*s* (Methods 4.10). Interestingly, for models with receptor recycling from VEE, the VEE recycling rate is much higher than the EE recycling rate when there is cAMP production in all the three compartments (Models 3.3 and 3.5, Appendix Table S3), in agreement with previously reported dynamics for the LHR (Jean-Alphonse et al. 2014; Sposini et al. 2017). Finally, irreversible receptor desensitization occurs mostly at the plasma membrane (with a probability of 0.01 for receptors to be internalized in endosomes) while most receptors within the endosomes are recycled back to the plasma membrane.

Given relatively comparable receptor recycling and ligand unbinding rates, but markedly contrasted signaling rates between the plasma membrane and endosomal compartments, we expect - in light of Sect. 2.1 - that internalization inhibition will primarily reduce signaling efficacy. Simulating dose-response experiments with decreasing internalization rates (Methods 4.8) using our best models (3.1 and 3.2), we show in Fig. 7 that the model indeed predicts that internalization inhibition drastically reduces signaling efficacy, with no apparent effect on potency. Similarly, biasing the spatial distribution of receptors - by inducing ligand-receptor accumulation in the third compartment - has virtually no effect on potency but substantially impacts efficacy. However, the large parameter uncertainty and the lack of agreement between the two best models complicate interpretation: for model 3.1, efficacy decreases as the proportion of *LR*_3_ increases, whereas in model 3.2, efficacy increases as the proportion of *LR*_3_ increases. These contrasting predictions highlight that experimental approaches capable of biasing receptor spatial distribution will be required, both for discriminating between models and for reducing parameter uncertainties within a given model.

**Figure 7.**
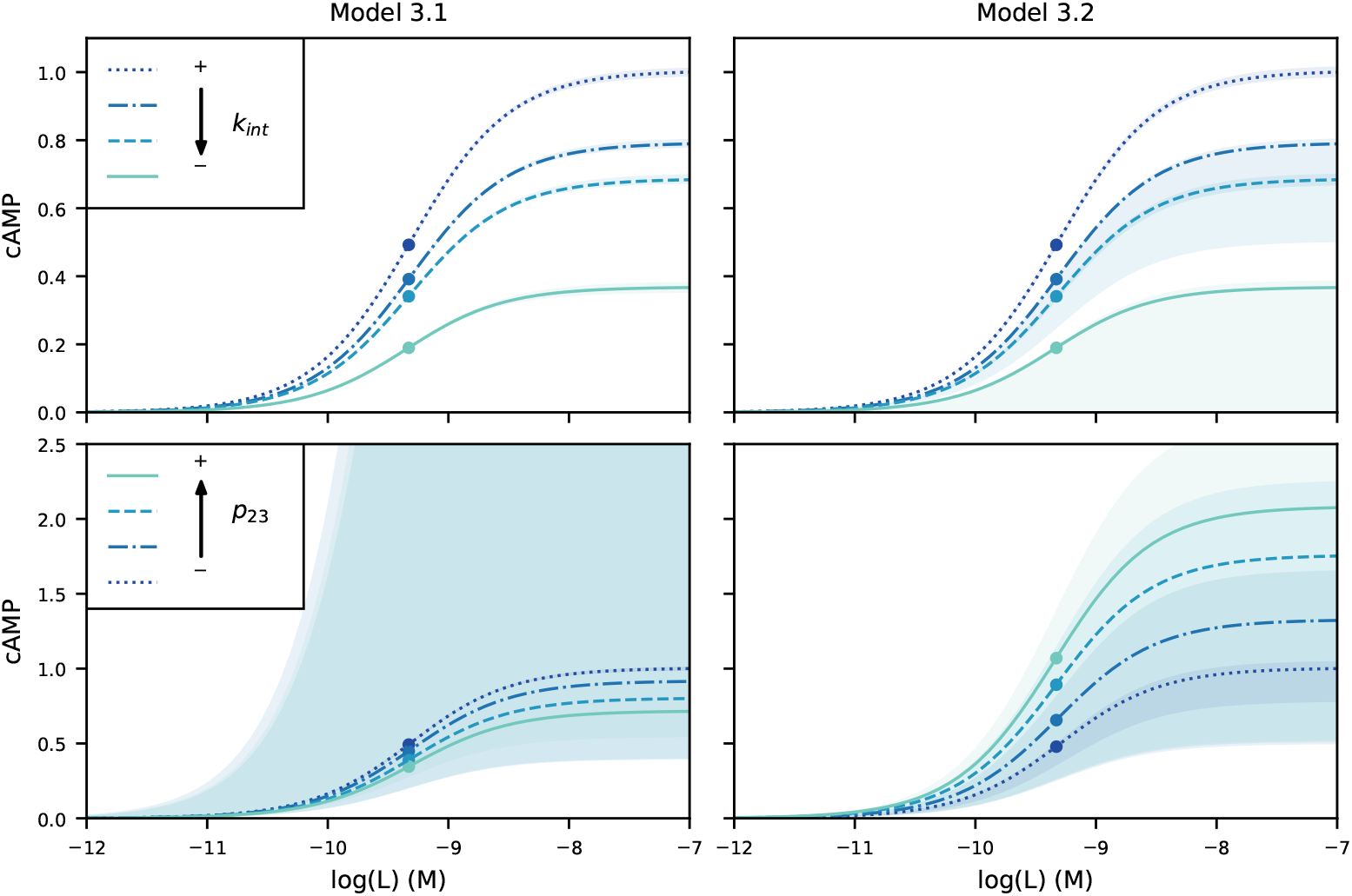
Dose response prediction of the total quantity of cAMP with internalization inhibition and with biased receptor spatial distribution for models 3.1 and 3.2 with 95% profile-based confidence intervals. The different values of internalization are by variying the parameters *ps*_2_: *ps*_2_ = 1, (dotted line), 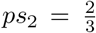 (dash-dot line), 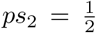 (dashed line), no internalization: *ps*_2_ = 0 (solid line). The different values for *p*_23_ are 0 (dotted line), 0.3 (dash-dot line), 0.7 (dashed line) and 1 (solid line) while the internalization rate remains constant for each model (*k*_*int*_ = *k*_*lr*13_ + *k*_*lr*12_, *k*_*lr*13_ = *p*_23_*k*_*int*_ and *k*_*lr*12_ = (1 − *p*_23_)*k*_*int*_). Circles represent the value of *EC*_50_. The remaining parameters used for the prediction are those found for the eight models selected by the AIC (Appendix Table S3).

### 2.4 Model validation

We also performed additional experiments for model validation, thet were not used for initial model selection or fitting. The three BRET assays (cAMP-EPAC, *PM*_*R*_-CAAX, and *EE*_*R*_-FYVE) were used to monitor total cAMP levels, receptor localization at the plasma membrane, and ligand-receptor complexes in early endosomes (EE) respectively. The plasma membrane-localized CAAX sensor targets YPet to the plasma membrane, enabling measurement of surface receptor levels, whose signal decreases upon ligand-induced internalization. This approach was used in a pulse-chase paradigm as opposed to continuous stimulation. In this design, the ligand was washed out after some minutes of exposure (Fig. EV1-validation data). Pulse-chase experiments are particularly well suited for dissecting the molecular events that occur after ligand binding to its receptor at the plasma membrane, while preventing new ligand binding. We additionally designed an experiment to characterize FSH binding to its cognate receptor, using a BRET-based dose-response assay. This assay relies on a recombinant FSH fused to the fluorescent acceptor mNeonGreen (*FSH*^*mNG*^) and a FSHR with a N-terminal NanoLuciferase tag (*Nluc* − *FSHR*) as the BRET donor. We further performed a competitive binding experiment which allowed indirect quantification of reference FSH binding through the dose-dependent decrease in *FSH*^*mNG*^-dependent BRET signal.

#### 2.4.1 The best model is able to predict data with pulse-chase ligand stimulation

Figure 8 shows the predictive performance of these models in representing pulse-chase data for a single ligand concentration, across all observables (cAMP, *PM*_*R*_, and *EE*_*R*_)(Fig. EV5 for all ligand concentrations). Nearly all validation data points (≈ 92% for all the concentrations of ligand) fall within the 95% confidence interval of the statistical error model. Although a systematic model bias is apparent, this may be attributable to variability between independent experiments, as observed in Fig. 4. Nevertheless, one notable model limitation emerges in the pulse-chase condition: following ligand washout, the number of membrane receptors first decreases rapidly due to internalization, then rebounds after approximately 20 minutes, presumably reflecting the fraction of receptors recycled back to the plasma membrane (Fig. 8-right column). This biphasic dynamic is not captured by our model, possibly because the large experimental variability obscures the underlying trend - most data points nonetheless remain within the confidence interval derived from the statistical observation model.

**Figure 8.**
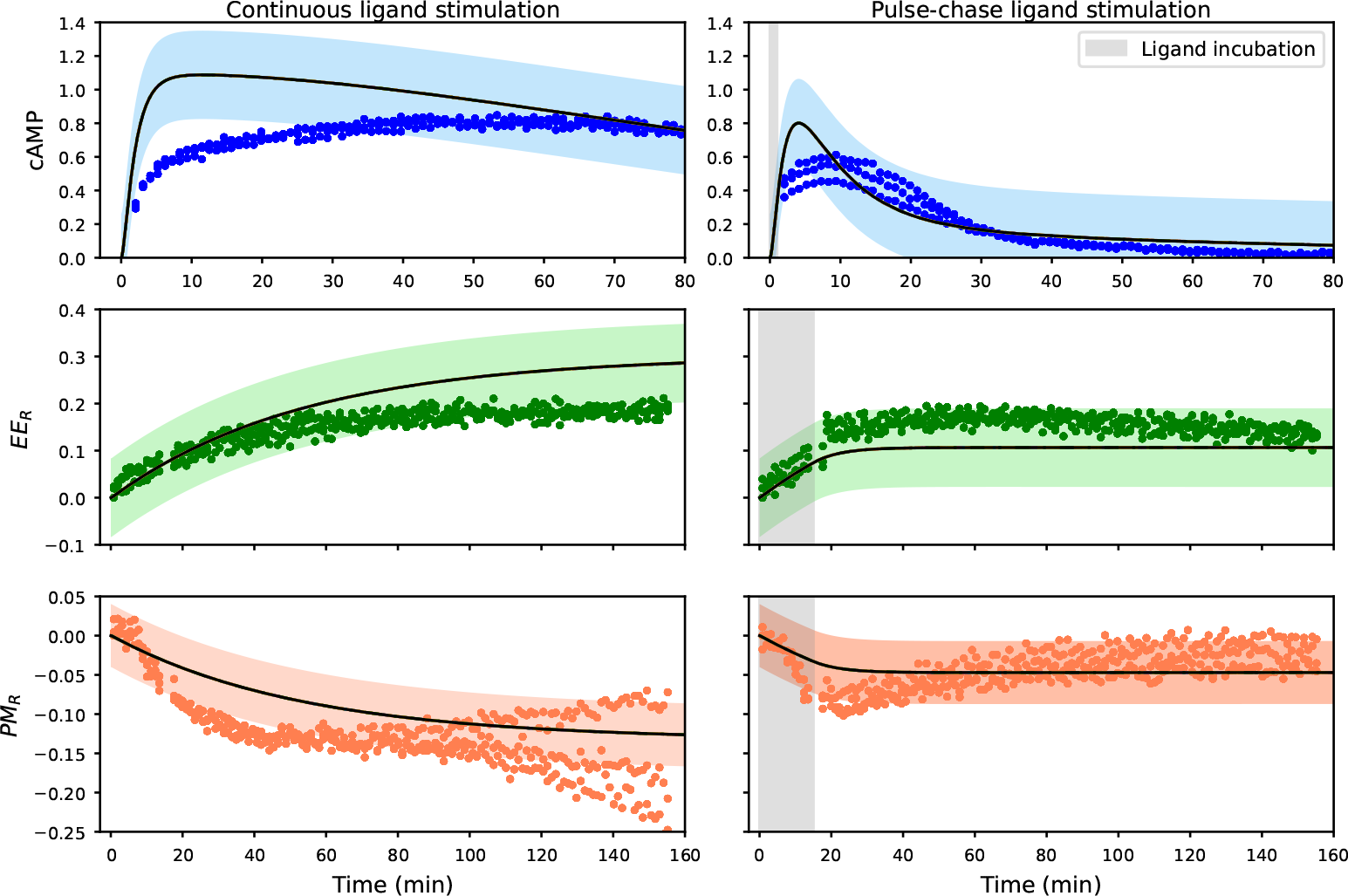
Prediction of observables with data validation for the 13 best models. The first column shows results under continuous ligand stimulation, and the second column under pulse-chase stimulation, both at a single FSH concentration of 30 nM. The first row displays cAMP levels, followed by *LR*_3_-FYVE (second row) and plasma membrane receptors-CAAX (third row). Scatter points represent validation data under both conditions (continuous and pulse-chase stimulation). Shaded areas indicate the 95% confidence interval of the statistical observation model across the 13 best models. The parameter values used for these predictions correspond to the best estimates obtained during model selection (Appendix Table S3).

#### 2.4.2 The best model correctly predicts binding affinity

To further validate our methodology, we verified the ligand binding affinity predicted by the model. The best model estimated the binding affinity of native FSH at *K*_*D*_ = 0.598 nM, with *K*_*D*_ comprised between (0.580, 0.622)nM, which is consistent in magnitude with previously reported values (Zariñn et al. 2020). To directly assess this prediction, we designed a dedicated competitive binding experiment. The concentration of *FSH*^*mNG*^ was held fixed while unlabelled FSH was added in excess (Fig. EV1, last row) to describe a competition experiment. This approach allowed indirect quantification of reference FSH binding through the dose-dependent decrease in BRET signal. The binding assay corresponds to the traditional equilibrium function with *f* the amount of ligand-receptor complexes:

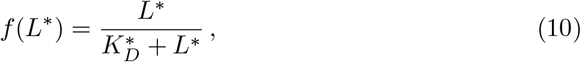

where *L*^∗^ is the fluorescent *FSH*^*mNG*^ concentration, and 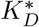 its binding affinity to FSHR, while the competitive assay corresponds to:

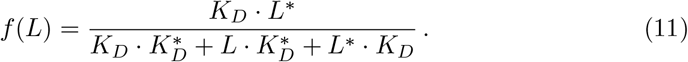

We jointly optimized Eqs. (10)-(11) in 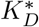, with *K*_*D*_ fixed to its model-prediction (*K*_*D*_ = 0.598 nM) with a least squares fitting. Given the good data-fitting in Fig. 9 and the fact that the fitted 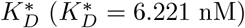 is very close to the *EC*_50_ measured in the experiment, we conclude that the model-predicted *K*_*D*_ value is coherent with our competitive binding assays.

**Figure 9.**
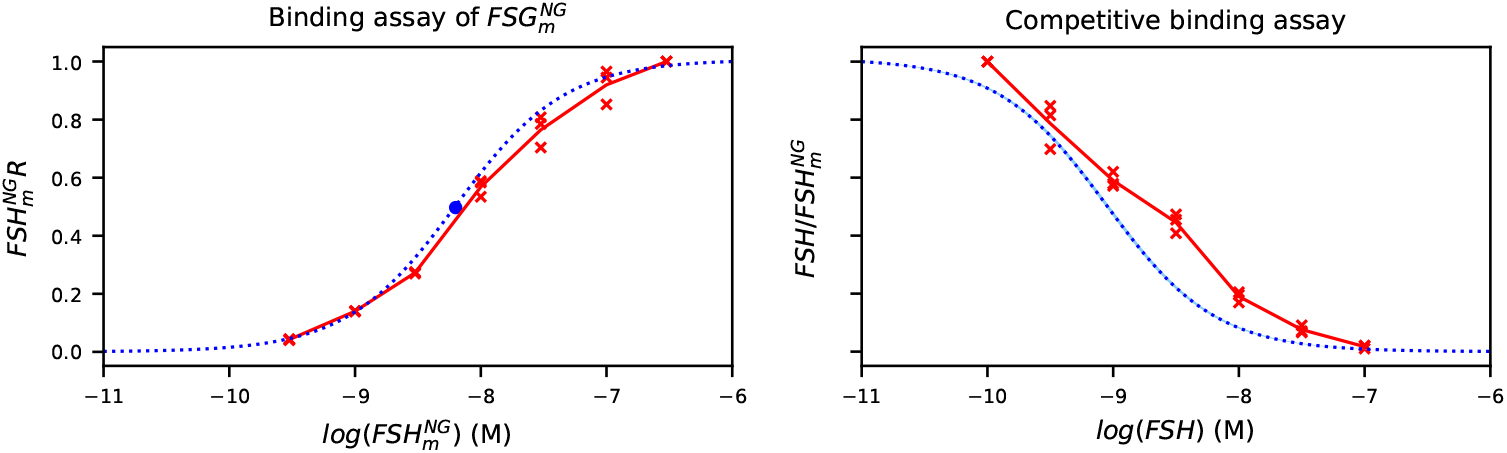
Validation of the FSH binding affinity. Binding assay of *FSH*^*mNG*^ to FSHR (left panel) and competitive binding of FSH to FSHR in presence of *FSH*^*mNG*^ (right panel, 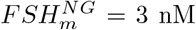). Data points are shown in red crosses, the solid red line represents the mean of the data and dotted blue line to the prediction. The fitted *FSH*^*mNG*^ binding affinity constant is shown by the circle on the left panel.

## 3 Discussion

The classical framework of GPCR signaling, in which receptor activation at the plasma membrane initiates downstream signaling cascades before being terminated through receptor internalization, has been substantially revised. For a growing number of GPCRs, internalization does not simply terminate signaling. Rather, receptors remain active within endosomal compartments, where they promote signaling responses that are qualitatively distinct from those initiated at the plasma membrane - a spatial organization that appears essential for appropriate physiological responses (Lyga et al. 2016; White et al. 2021; Gourdon et al. 2025). Nevertheless, many aspects of receptor trafficking remain poorly understood, and current experimental approaches face inherent limitations in disambiguating the respective contributions of plasma membrane and endosomal signaling to the overall cellular response. In this context, a mathematical modeling approach offers a valuable complementary framework. By deriving and analyzing theoretical results, such an approach can generalize the impact of receptor trafficking on cellular responses and bridge mechanistic assumptions and experimentally observable outputs. The aim of this modeling framework was to investigate the impact of receptor trafficking and subcellular receptor distribution on signaling responses, and more specifically to understand how receptor trafficking shapes the overall cellular response. Analysis of dose-response profiles provides a mean to characterize cellular responses across different receptor systems and to evaluate the effects of perturbations on the signaling output.

Several simplifying assumptions were made to ensure analytical tractability and interpretability of the dose-response relationships (Methods 4.1). First, the ligand was assumed to be in large excess relative to the receptor amount, such that its concentration can be treated as constant, yielding a linear ligand-receptor association rate. Second, ligand and receptor degradation were neglected, as including such terms would preclude the calcul of analytical expressions (null function). Third, the rate of second messenger degradation was assumed to be identical across compartments, despite potentially distinct production rates, in order to reduce the number of free parameters and improve interpretability. Fourth, all systems were initialized with the total receptor pool only, under the assumption that receptor occupancy at the initial time is negligible - although constitutive receptor activity is acknowledged. Finally, ligand-receptor dissociation within intracellular compartments was not considered. However, our results can be further generalised for (i) nonlinear cAMP production rate; (ii) compartment-dependent cAMP degradation rate; (iii) time-dependent ligand association kinetics (*L* + *R* ↔ *LR*).

Theoretical results allow the kinetic contribution of different parameters to be assessed by examining the shape of dose-response curves and their associated pharmacological descriptors, such as efficacy and potency. Our analysis establishes a link between model predictions and experimental observations, and reveals counter-intuitive effects of receptor trafficking and spatial distribution on the observed *EC*_50_ (Fig. 1). When studying the impact of receptor trafficking on the signaling response, researchers routinely disrupt dynamin or clathrin activity through the use of drugs such as Dyngo4a and PitStop2 (Gourdon et al. 2025), or via overexpressing a dominant negative mutant protein like DynK44E (Blythe et al. 2025). Recently, it has been shown that inhibiting endocytosis by overexpressing DynK44E leads to a significant decrease in ligand-induced cAMP response for *β*2-adrenergic receptor (*β*2AR) and vasoactive intestinal peptide receptor 1 (VIPR1) (Blythe et al. 2025). Further, this strategy had no effect on potency for these receptors. In the light of our results (Subsect 2.1.1), this suggests that for *β*2AR and VIPR1, the endocytic response is strong 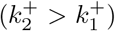, and ligand dissociation and receptor recycling occur on a similar timescale (*k*_*off*_ ≈ *k*_*rec*_) (Fig. 1). By constrast, for the luteinizing hormone receptor (LHR), inhibiting endocytosis with chemical compounds has been reported to induce a strong increase of the observed potency, with no or moderate impact on the efficacy for both hCG- and LH-induced cAMP response (“shift to the right”, Fig. 1)(Gourdon et al. 2025). This suggests that for the LHR, the compartmentalized cAMP production is balanced 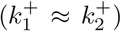, but receptor recycling is slower than ligand unbinding (*k*_*off*_ *> k*_*rec*_).

Several authors have shown that LHR is internalized into different types of endosomes (*e*.*g*. VEE and EE), with different downstream signaling efficiency and recycling kinetics (Sposini et al. 2017; Jean-Alphonse et al. 2014). In Raynaud et al. (2025), the authors designed an intracellular antibody fragment (i.e. an intracellular VHH), which drives receptor accumulation in EEs at the expense of VEEs (increased *LR*_3_ at the expense of *LR*_2_) associated to a decreased of overall receptor recycling. Interestingly, this innovative tool had no effect on receptor internalization, thus providing a way to finely tune ligand-induced receptor endosomal distribution. Receptor accumulation in EEs was associated with a reduction in global cAMP production. This observation suggests that signaling from EEs is less efficient than that originating from VEEs or the plasma membrane - consistent with a decrease in *E*_*max*_ (condition (18), Fig 2c/d). The fact that the recycling rate from EEs is presumably lower than the one from VEEs (*k*_*r*21_ *> k*_*r*31_) suggests a potential shift to the left (decrease of *EC*_50_), which constitutes an experimentally testable prediction for future model validation (Fig 2a/c). More broadly, condition 18 - which integrates trafficking and signaling production parameters to determine the conditions under which *EC*_50_ increases - underscores the complexity of GPCR signaling when three spatially distinct pools of active ligand-receptor complexes coexist. This complexity renders straight-forward biological interpretation challenging, and calls for the joint use of dedicated mathematical modeling and carefully designed experimental approaches to rigorously disentangle the interrelationships between receptor trafficking kinetics and signaling response dynamics.

The parameter estimation framework developed for the FSHR case study allowed us to determine the order of magnitude of most kinetic parameters, which were, to the best of our knowledge, previously unknown (Sect. 2.2). In particular, the joint analysis of kinetic dose-response experiments - encompassing receptor internalization kinetics under both control and perturbed conditions - together with an exhaustive model selection procedure, supports the hypothesis that the FSHR is actively signaling from the plasma membrane and two distinct endosomal compartments, or alternatively from the plasma membrane and a single endosomal compartment consistent with early endosomes. These conclusions are in line with earlier biological studies (Sposini et al. 2017; Jean-Alphonse et al. 2014), although the scenario involving signaling only from early endosomes and plasma membrane warrants careful interpretation. Indeed, existing evidence tends to suggest that very early endosomes constitute the primary active intracellular signaling compartment, even if there is no definitive experimental confirmation to date. Better understanding spatial dynamics of gonado-induced response could help improve our knowledge of the complex signaling networks it regulates, and therefore improve the design of medical treatments for reproductive disorders.

At present, it is not possible to determine a unique model, but rather a cluster of plausible models. However, a first way to distinguish whether receptors remain active in all compartments or only in the PM and EE is to analyze predictions as the number of EE increases (Fig. 7). Assuming activity in all three compartments leads to a decrease in efficacy, consistent with Raynaud et al. (2025), supporting the idea that receptor activity is not restricted to only two compartments. In comparison, this result could validate results found in Subsect. 2.1.2. In fact, the parameter range of Model 3.1, which leads to a decrease in efficacy due to receptor accumulation in the third compartment (Fig. 7), confirms condition (18): the number of active receptors is higher for *LR*_1_ and *LR*_2_ than from *LR*_3_ (Appendix Table S3). Interestingly, an in-depth parameter identifiability analysis across this cluster of models revealed that substantial intracellular signaling can occur despite a small number of internalized receptors, a configuration arising from slow internalization combined with fast recycling rates. These predictions are biologically plausible, although experimentally validating them remains challenging, as precise measurements of receptor internalization and recycling kinetics are difficult to obtain. Some studies have partially addressed this by characterizing receptor trafficking and quantifying the number of receptor-positive endosomes labeled with specific compartment markers - such as APPL1 for very early endosomes (*LR*_2_/VEE) and EEA1 for early endosomes (*LR*_3_/EE). In Sposini et al. (2020), for the FSHR, 42% of receptor-positive endosomes are positive to APPL1 and 36% of receptor-positive endosomes are positive to EEA1. Due to the high model uncertainty, direct comparison of proportions is not possible, but our results indicate that a similar order of magnitude of *LR*_2_/VEE and *LR*_3_/EE is coherent with the model (confidence interval cross each others in Table 1). The fact that models without recycling from EE are desensitised is particularly interesting in the case of LHR, since the first assumption is that receptors from EE are not recycled. One explanation for the large differences predicted by the model in signalling efficiency between the internal and plasma membrane compartments could be that cAMP degradation is assumed to be identical in all compartments. The proportion of internalized receptors might be drastically higher if local cAMP degradation were taken into account in the model.

In addition, Sposini et al. (2017) reported that receptor recycling is rapid, peaking at approximately 5 min before reaching a plateau - a finding consistent with the time scales inferred in our model (Table 1). The validity of these inferred time scales is further supported by the ability of the model to predict cAMP kinetics under ligand pulse-chase validation experiments as well as ligand binding kinetics. Among the best-fitting models, the main differences lie in the quantity of ligand-receptor complexes within intracellular compartments and in the magnitude of internalization and recycling fluxes. Further model validation could be achieved through experimental measurement of internalization and recycling kinetics via single-particle tracking, as well as quantification of receptor proportions in VEEs and EEs by whole-cell microscopy imaging. Additionally, compartmentalized cAMP biosensors - which could be designed once distinct endosomal compartments are more precisely characterized using specific molecular markers - could provide an indirect means of discriminating between competing trafficking hypotheses, particularly with respect to VEEs.

The trafficking of GPCRs was modeled using ordinary differential equation systems describing the temporal evolution of the relevant molecular species. This modelling approach improves our understanding of how cells encode receptor signaling by providing a mechanistic interpretation of underlying biological processes, and can readily be extended to other GPCRs by adjusting trafficking dynamics and the number of compartments, and implementing the corresponding biological hypotheses. To ensure analytical tractability, several simplifying biological assumptions were made to reduce the number of chemical species, despite the known complexity of the system. This parsimony represents both a strength and a limitation of the model: while it yields an interpretable and generalizable framework, it inevitably omits certain biological interactions. Several natural extensions could be envisioned to progressively relax these assumptions. At the level of receptor-G protein coupling, the G protein cycle could be incorporated through a ternary complex formalism (Chen 2003), and further enriched by accounting for the subcellular trafficking of G proteins themselves - recent evidence highlighting the role of *G*_*αs*_ trafficking in intracellular signaling (Sokrat et al. 2024; Gourdon et al. 2025) makes this a particularly relevant extension. The role of *β*-arrestins, whose involvement in receptor internalization has recently been substantially revisited, also warrants explicit consideration (Irannejad and von Zastrow 2014; Eichel and von Zastrow 2018; Sposini and Hanyaloglu 2018; Eiger et al. 2023; Daly et al. 2023). Finally, the framework could be extended to additional subcellular compartments, irrespective of whether signaling responses are produced therein (Crilly and Puthenveedu 2021).

We employed simplified, or ‘toy’, models of receptor signaling cascades to isolate and precisely characterize the specific contribution of receptor trafficking to the overall cellular response. Since the introduction of the operational model, numerous mathematical frameworks have been developed to capture the complexity of GPCR-induced signaling, addressing aspects such as constitutive activity, G protein cycling, and the activation of multiple effector pathways (Chen 2003; Bridge 2010; Woodroffe et al. 2010; Bridge et al. 2018; Finlay et al. 2020; Carvalho et al. 2021; Bridge et al. 2024). Yet, to our knowledge, few modeling studies have explicitly focused on receptor trafficking, and existing frameworks may now require re-evaluation in light of the recognized importance of signaling compartmentalization. Beyond receptor trafficking itself, a comprehensive understanding of compartmentalized GPCR signaling requires accounting for the coordinated dynamics of multiple cellular actors. These include the differential involvement of *β*-arrestins in receptor sorting (Heitzler et al. 2012; Tóth et al. 2023; Liu et al. 2025), and the diffusion of second messengers such as cAMP (Anton et al. 2022). Although the modeling of these phenomena remains in its early stages, their integration into a unified framework will be essential for a fully mechanistic description of spatially organized signaling.

Finally, the present framework relies on deterministic ODE modeling, which captures the average behavior of a cell population. Kinetic dose-response profiles should therefore be interpreted as mean cellular responses across a population, rather than as descriptions of individual cell behavior. With the rapid advances in single-cell imaging and single-cell resolution measurements of signaling pathways, the development of stochastic models represents a promising extension of this work, offering a principled framework for characterizing cell-to-cell variability in signaling responses (Duso and Zechner 2020; Kolbe et al. 2022).

## 4 Methods

### 4.1 Biological assumptions for modeling

For all the models 1, 2, 3 general assumptions were:

1. the quantity of ligand (*L*) was considered to be in excess compared to the total quantity of receptors and was therefore taken as a constant, resulting in ligand kinetics that are independent of time;
2. ligand and receptor degradations were ignored;
3. the second messenger (cAMP) was produced in each compartment proportionally to the ligand-receptor complexes, but was degraded linearly independently of the compartment type 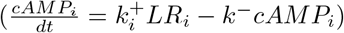;
4. we started with an initial quantity of free receptors at the plasma membrane (*R*_1_(0) = *R*_0_) and the initial quantity of all other species was null (initial points of the ODE is wrote *x*(0));
5. there was no dissociation of the ligand-receptor complexes in the intracellular compartments and the complex dissociated upon recycling to the plasma membrane.

In addition, ligand-receptor complexes degradation was ignored in 1, 2.

### 4.2 Mathematical modeling

All the models in our study were studied using the ordinary differential equation formalism, following the evolution concentration of species over time. Then, the equilibrium of each system was calculated to obtain to study the influence of each parameters on the dose-response.

#### 4.2.1 Model 1

Model 1 studied in the Subsect. 2.1.1 leaded to the following ODE system:

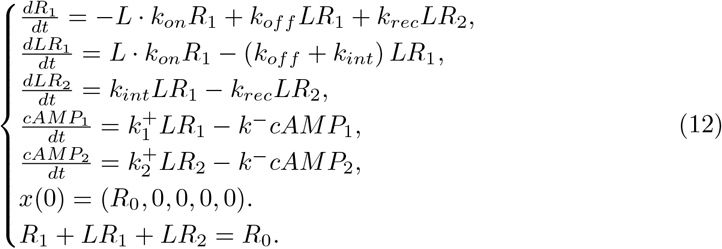

In short, through binding (resp. unbinding) reactions at rate *k*_*on*_ (resp. *k*_*off*_), lead to the formation of a ligand-receptor complex (*LR*_1_) at the plasma membrane. This complex is then internalized into the second compartment at rate *k*_*int*_, while the receptor is recycled back to the membrane at rate *k*_*rec*_. The second messenger (cAMP) is produced in each compartment, in proportion to the amount of ligand-receptor complex *LR*_*i*_, with *i* = 1, 2 with compartment-specific production rates 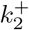 at the plasma membrane and 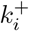 in the intracellular compartment and is degraded according to first-order kinetics, at a rate *k*^−^. Appendix Table S1 summarizes all variables and parameters of this model.

By mass conservation, the receptor trafficking subsystem (*R*_1_, *LR*_1_, *LR*_2_) can be reduced to a two-dimensional affine system whose system matrix is stable (by simple calculations). Thus, the receptor variables are globally exponentially stable, and the same holds for the second messenger cAMP, due to its first-order degradation kinetics. To compute the steady state, consider the system:

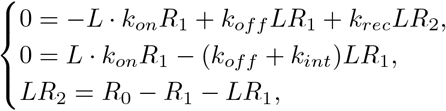

we deduce thanks to simple algebraic calculations the expressions for *LR*_*i*_:

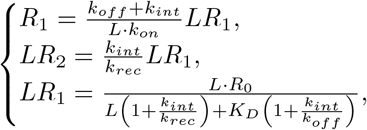

and then using:

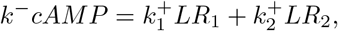

we obtain Eq. (1). The later Eq. (1) has a similar expression as the standard phenomenological dose-response curve which we recall:

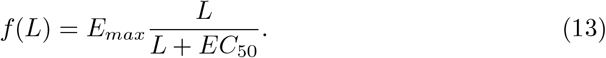

In Eq. (13), *E*_*max*_ quantifies the ligand efficacy (maximum of the cellular response) and *EC*_50_ is the ligand potency (quantity of ligand needed to obtain half of the maximum of the dose response) (Black 1983). Identifying *E*_*max*_ and *EC*_50_ in Eq. (1) leads to Eqs. (2)-(3).

#### 4.2.2 Model 2

Model 2 studied in Subsect. 2.1.2 leads to the following ODE system (Eq. 14):

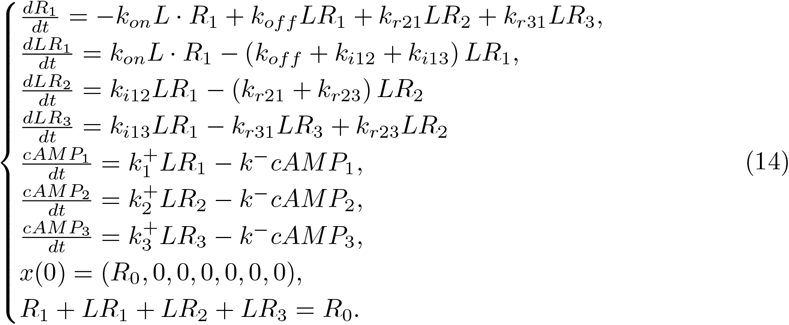

In short, binding and unbinding reactions, occurring at rates *k*_*on*_ and *k*_*off*_ respectively, lead to the formation of a ligand-receptor complex (*LR*_1_) at the plasma membrane. The ligand-receptor complex *LR*_1_ is then internalized in two intracellular compartments, called *LR*_2_ or *LR*_3_, at rates *k*_*i*12_ and *k*_*i*13_, respectively. *LR*_2_ is recycled at rate *k*_*r*21_ (respectively *LR*_3_ at rate *k*_*r*31_) and receptors in the second compartment, called *LR*_2_, can transfer to the third compartment, called *LR*_3_, at rate *k*_*r*23_. The second messenger (cAMP) is produced in each compartment proportionnally to the ligand-receptor complex *LR*_*i*_ with compartment-specific production rates 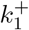 for *i* = 1, 2, 3, and is degraded according to first-order kinetics, at a rate *k*^−^. Appendix Table S1 summarizes all variables and parameters of this model. Using the same methodology as in Subsection 4.2.1, we derive the analytical formula for the steady-state cAMP response, which reads as follows. By simple algebraic calculations, we obtain the following expression for *LR*_*i*_:

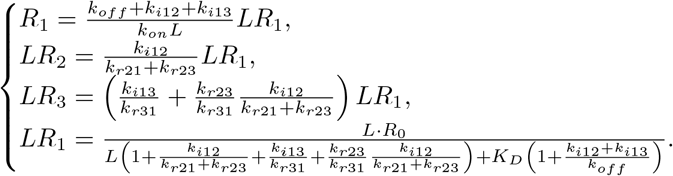

The steady-state for cAMP is the following:

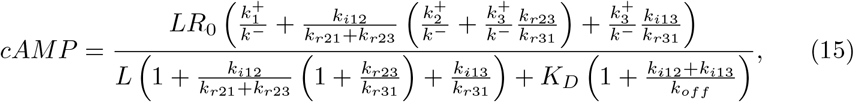

with the values of potency and efficacy:

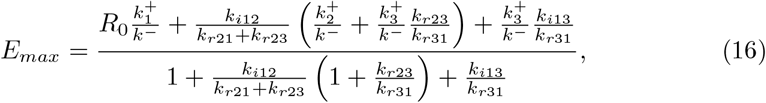

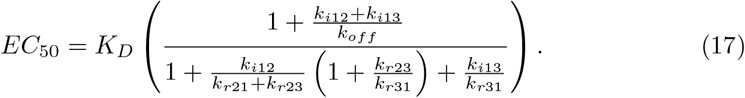

Now using *k*_*i*12_ = (1 − *p*_23_)*k*_*int*_ and *k*_*i*13_ = *p*_23_*k*_*int*_, we obtain efficacy, potency and transducer ratio values given in Eqs. (6)-(8). We study the effect of increasing the proportion of receptors in the third compartment (*p*_23_) on the efficacy *E*_*max*_, potency (*EC*_50_) and *τ* (Eqs. (6)-(8)), to obtain the following necessary and sufficient conditions: *EC*_50_ is increasing according to *p*_23_

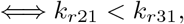

*E*_*max*_ is increasing according to *p*_23_

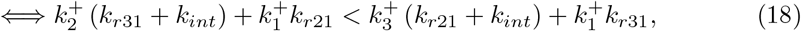

and, *τ* is increasing according to *p*_23_

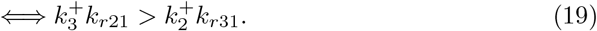

#### 4.2.3 Model 3

The general model 3 for the parameter estimation problem (Sect. 2.2) is the following ODE system (Eq. 20 and Appendix Table S1):

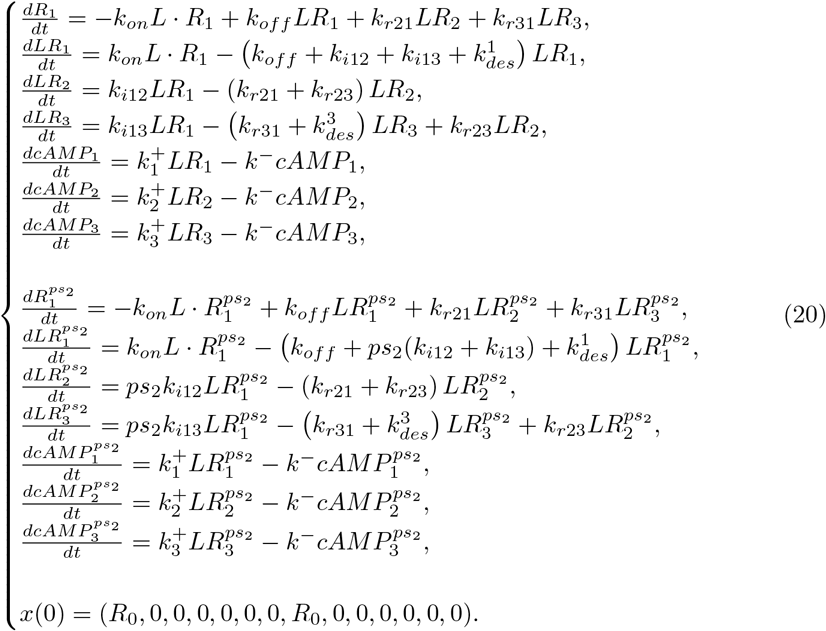

This model is exactly the same model as Eq. (14) but includes the desensitization/degradation of *LR*_1_ at the plasma membrane 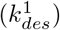 and of *LR*_3_ in the early endosomes 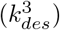. Appendix Table S1 summarizes all variables and parameters of this model.

### 4.3 Reparameterization of the equations for the parameter estimation problem

The parameter estimation problem includes different types of parameters, such as initial conditions, kinetic parameters, error parameters and data parameters, totalling 21 parameters (Appendix Table S2).

To decrease the number of parameters, the first step is to reparameterized the model. All species quantity are divided by *R*_0_ and *k*_*off*_ gives the time scale. We note *k*^∗^ as the normalized parameters: 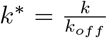. In addition, production parameters can be linked together by following this method. Let 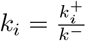, for *i* = 1, 2, 3, then equation is obtained,

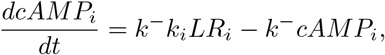

and we note *EPAC* = *α*_1_ ∑ *cAMP*_*i*_, with *i* the number of the compartments’ type. With variables changes, a reparameterized system with less parameters is obtained:

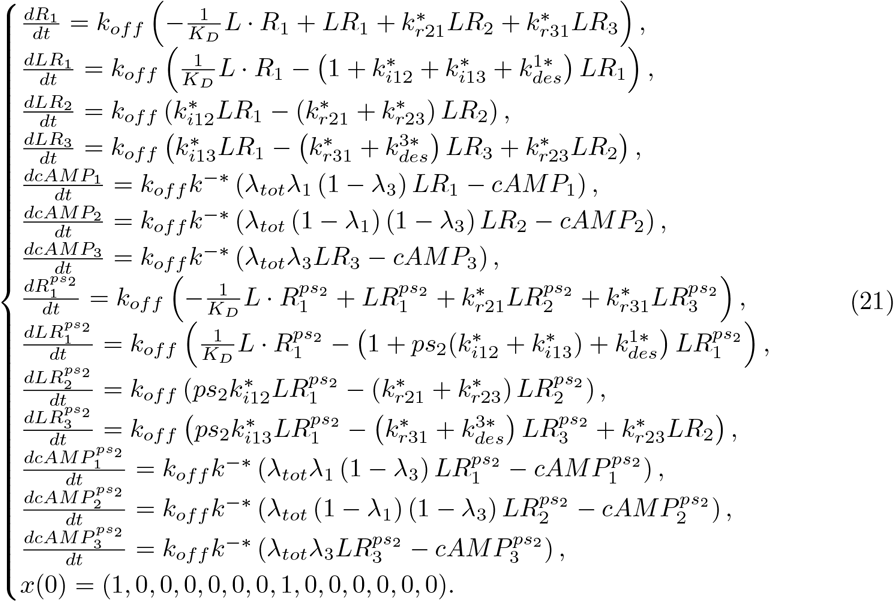

with:

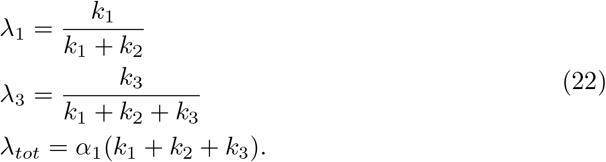

The observables-equation reparameterized system is

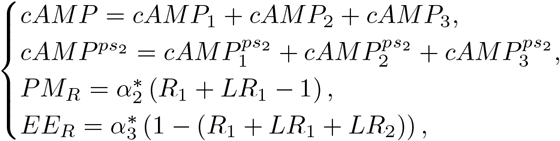

with 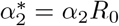 and 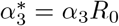.

The number of parameters decreases from 21 to 19, by deleting the initial quantity of receptor, *R*_0_, and by replacing, 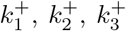 and *α*_1_ by *λ*_1_, *λ*_3_ and *λ*_*tot*_ (22). The number of parameters reparameterized is summarized in table 3.

## 4.4 Parameter estimation

The parameter estimation problem is implemented using the Python-based tools PEtab (Schmiester et al. 2021) and pyPESTO (Schälte et al. 2023). PEtab provides a structured framework for parameter estimation by combining several tables, including the network, measurements, conditions, parameters, observables, and visualization results, thereby linking them consistently. This organization makes it easier to modify model parameters rapidly and to associate different measurements with observables under identical biological conditions. pyPESTO is then used to solve the PEtab problem and provides useful tools for analyzing the results. In particular, it evaluates the log-likelihood function for each parameter vector *θ*, together with the noise parameter the vector of parameters *σ*_*i*_ associated for each observable (Eq. 9). We assume Gaussian noise for each observable. Let *y*_*ij*_ denote the *j*-th observable at time *t*_*i*_, and let *f*_*j*_(*t*_*i*_; *θ*) be the corresponding ODE solution. The negative log-likelihood is

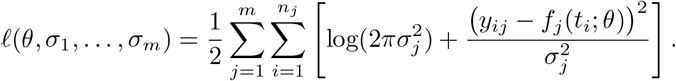

To resolve the parameter estimation, the algorithm was run 1000 times to ensure a satisfactory fit with the optimizer Fides. Several runs reach to the same optimum for the 13 best models leading to algorithm convergence (Appendix Figure S 1).

### 4.5 Model selection

Our goal is to identify a parsimonious model that achieves robust parameter estimation and addresses key hypotheses. We performed model selection using Petab and pyPESTO, which employ deterministic gradient descent with multi-start optimization to compute the Akaike Information Criterion (*AIC* = 2*k* − *ln*(*L*)), where *k* is the number of parameters and *L* is the likelihood estimate. This criterion penalizes overly complex models. To resolve the model selection, the algorithm was run 1000 times on each model and the best AIC from the 1000 runs was selected. To validate the selected model, we applied the Δ_*AIC*_ criterion (Burnham and Anderson), which compares each model’s AIC to that of the best models. Δ_*AIC*_ *<* 2 are considered as likely as the best model; 2 *<* Δ_*AIC*_ *<* 7, as suitable alternatives; 7 *<* Δ_*AIC*_ *<* 10, less relevant; and those with Δ_*AIC*_ *>* 10 are rejected.

### 4.6 Structural and practical identifiability

To verify the theoretical stability of the complete model after reparameterization (Eq. (21)), we employed StructuralIdentifiability (Hong et al. 2019, 2020; Dong et al. 2023). This tool allows us to assess the structural identifiability of the model independently of biological data. Indeed, based on our experience, not all model variables are experimentally accessible, even though these species and their associated parameters are essential for the model. To evaluate practical identifiability relative to available biological data, we use the Profile Likelihood Estimate (PLE) method (Kreutz et al. 2013):

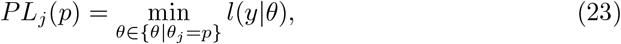

where the negative log-likelihood is evaluated as a function of the values *p* of a parameter component *θ*_*j*_, while all other parameters *θ*_*i*_, *i* ≠ *j* are reoptimized. The values comprise in the 95% confidence interval are kept. The profile likelihood estimate for models 1 and 2 is shown in Appendix Figure S2.

### 4.7 Error

Throughout this paper, two types of uncertainties are presented to characterize the data: the statistical error and the profile biaised error. The statistical error captures the link between the observables and the data. It corresponds to the parameters *σ*_1_, *σ*_2_ and *σ*_3_, associated respectively with observables (Eq. (9)). The parameter estimation problem infers the variance of the data with respect to each observable. Consequently, when plotting the observables, it is essential to include the statistical error as follows:

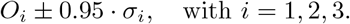

The profile biaised 95 % confidence interval accounts for the uncertainty in the predictions. Since the parameters are each associated with a confidence interval estimated via the Profile Likelihood Estimation (PLE) (Eq. 23), they do not take a single unique value. It is therefore essential to propagate these uncertainties when computing predictions for quantities other than the observables. The predictions are computed for all parameter values within the confidence interval, and the minimum and maximum values of each prediction are retained to define the uncertainty bounds.

### 4.8 Dose-response experiments

Since Eq. (20) includes irreversible receptor desensitization, all species have a steadystate value of 0, unlike the models considered in Sect. 2.1. Therefore, rather than focusing on the steady state, we calculated dose-response experiments and quantified the response by the area under the curve (AUC). We simulated Eq. (20) over one hour for different internalization rates in order to generate the dose-response profiles shown in Fig. 7.

### 4.9 Estimation of the constant affinity

The least squares algorithm from scipy.optimize is used to fit models (10)-(11) to the validation binding experiments, in order to estimate the value of 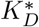.

### 4.10 Times, probability and proportion

#### 4.10.1 General formula

Endocytosis and recycling events are not almost certain events from a probability point of view given that the model includes irreversible receptor desensitization. Therefore, we calculated conditional expectation times using the following method (Kampen 2007). Given a continuous-time Markov chain (CTMC) of infinitesimal generator *L* on a state-space Ω ∪ *A* ∪ *B* where *A* and *B* are two disjoint absorbing ensembles, we calculated the solution of

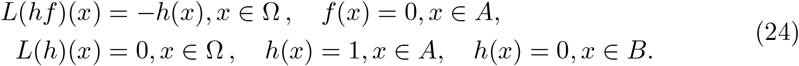

In Eq. (24), *h*(*x*) is the absorbing probability in *A*, starting from *x* ∈ Ω, and *f* is the conditional expectation time:

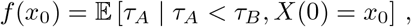

with *τ*_*A*_ and *τ*_*B*_ the absorbing times in *A* et *B* respectively.

#### 4.10.2 Endocytosis time and probability

We used the CTMC whose graph is the following:

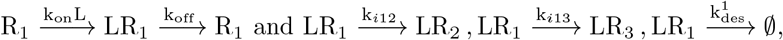

with Ω = {*R*_1_, *LR*_1_}, *A* = {*LR*_2_, *LR*_3_} and *B* = {∅} (the absorbing desensitization state). We obtained for the conditional expectation time for endocytosis, with *x*_0_ = *R*_1_:

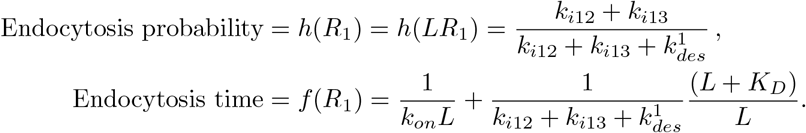

#### 4.10.3 Recycling time and probability

We used the CTMC whose graph is the following:

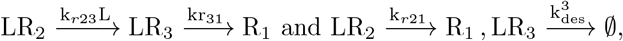

We set Ω = {*LR*_2_, *LR*_3_}, *A* = {*R*_1_}, and *B* = {∅} (the absorbing desensitization state). We obtained for the endocytosis, with *x*_0_ = *LR*_2_ and *x*_0_ = *LR*_3_:

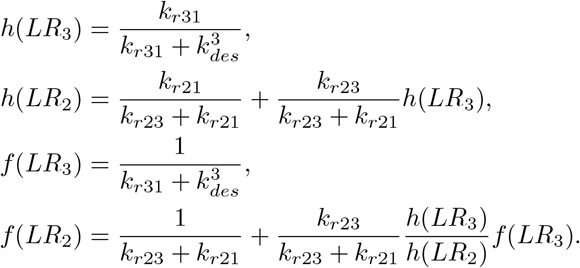

Finally, we averaged out these quantities according to the initial point, which is *LR*_2_ with probability 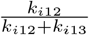, and *LR*_3_ with probability 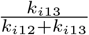. We thus obtained

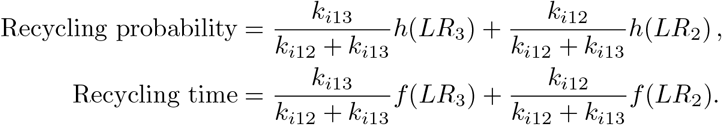

#### 4.10.4 Receptors proportion

The proportion of receptors is:

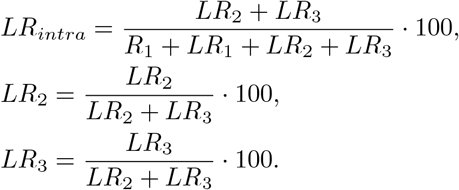

### 4.11 Biological data

All measurements were performed across 10 independent experiments per data type, revealing inter-experiment variability, likely attributable to differences in receptor and/or BRET sensor expression levels.

#### 4.11.1 Ligands and drugs

Recombinant FSH (GONAL-f^*R*^) was kindly provided by Merck and resuspended in mQ H_2_O.

FSH-mNeonGreen was designed in our group and produced in ExpiCHO-S^*TM*^ cells expression system. ExpiCHO-S^*TM*^ cells at 6.10^6^ living cells/mL density were transfected with 20 *µ*g of pcDNA3.1 encoding the FSH-mNG, according to ExpiCHO-S^*TM*^ Expression System (Gibco^*TM*^) manufacturer’s instructions. Cells were cultured for one day at 37^°^C/8% CO_2_ on a shaker platform (120 rpm) before supplementation with feed and enhancer solution (provided in the ExpiCHO-S^*TM*^ expression system transfection kit). Cells were then cultured for 12 days at 32^°^C/5% CO_2_ at 120 rpm. The culture was centrifuged at 500 g/30 min/4^°^C, and the supernatant was then collected. The supernatant was then centrifuged again at 5000g/30min/4^°^C. The newly collected supernatant was dialyzed in regenerated cellulose membrane tubing with a 6-8 kDa molecular weight cut-off (MWCO) (Spectra/*Por*^*M*^) against a pH 8.0 buffer (50 mM Tris HCl, 100 mM NaCl) overnight at 4^°^C under continuous stirring. The dialyzed supernatant was centrifuged (5000g/30min/4^°^C) to remove insoluble debris, and passed through a buffer equilibrated (pH 8.0, 50mM TrisHCl, 100mM NaCl) Protino^@^ Ni-IDA packed column (Macherey-Nagel^*TM*^). The column was washed with three times its volume of the following buffers: i) pH 8.0, 50mM TrisHCl, 100mM NaCl, ii) pH 8.0, 50mM TrisHCl, 1M NaCl, iii) pH 8.0, 50mM TrisHCl, 100mM NaCl. The FSH-mNG elution was performed with a pH 8.0, 50mM TrisHCl, 100mM NaCl, 500mM imidazole buffer. The FSH-mNG was buffer exchanged into a pH 8.0, 50mM TrisHCl, 100mM NaCl buffer, using 10DG Desalting Prepacked Gravity Flow Columns (BioRad Laboratories). The FSH-mNG was then concentrated using 30kDa MWCO Amicon Ultra centrifugation unit (Sigma-Aldrich) according to the manufacturer’s instructions. After analysis by Coomassie blue SDS-PAGE gel staining, purified FSH-mNG was stored at -20^°^C.

PitStop2 was purchased at Sigma-Aldrich, and used at a final concentration of 30*µ*M.

#### 4.11.2 Plasmids

The FSHR-RLuc8 plasmid was kindly provided by Pr. Aylin Hanyaloglu (Imperial College London, United Kingdom). All other plasmids were designed in our group and synthesized by Twist Bioscience. The cAMP BRET sensor NLuc-EpacD602A-VV-NES was designed from the cAMP BRET sensor NLuc-Epac-VV (Masuho et al. 2015) by adding a nuclear exclusion signal (NES) sequence and the D602A mutation position to improve the sensor’s dynamic range (Klarenbeek et al. 2015).

#### 4.11.3 Cell culture

Human Embryonic Kidney 293 (HEK293A) (Thermo Fisher Scientific) cells were cultured in DMEM (Eurobio) medium containing Glutabio and NaHCO_3_ and supplemented with 10% heat inactivated fetal bovine serum (FBS) (Eurobio), 100 IU/mL penicillin and 0.1 mg/mL streptomycin (Eurobio). Cells were kept at 37^°^C in a humidified 5% CO_2_ incubator.

#### 4.11.4 Bioluminescence Resonance Energy Transfer (BRET)

Forty thousand HEK293A cells per well were seeded in previously 0.01% poly-lysine coated 96-well plates, and transiently transfected in suspension using Metafectene Pro transfection reagent (Biontex Laboratories) according to the manufacturer’s instructions. DNA quantities used in each type of experiment are detailed in the following table:

| Type of experiment | Quantity of plasmid DNA transfected/well |
| --- | --- |
| cAMP measurement | FSHR: 25ng<br>cAMP sensor NLuc-EpacD602A-VV-NES: 20ng |
| Receptor trafficking | FSHR-RLuc8: 90ng<br>Lyn-Ypet, Ypet-CAAX or Ypet-FYVE: 20ng |
| Binding of FSH-mNeonGreen | NLuc-FSHR: 30ng |

48 hours after transfection, BRET measurements were performed upon addition of 5 *µ*M coelenterazine-H (Interchim) diluted in Ca^2+^/Mg^2+^free PBS, containing no or different concentrations of FSH. For experiments performed in presence of PitStop2, cells were pre-incubated 35 minutes in presence of drug-containing buffer before measurements and ligand stimulation. Signals were recorded for at least 60 minutes with a Mithras LB 943 plate reader (Berthold Technologies GmbH & Co.). BRET ratios were calculated as follows: 480nm/540nm for cAMP experiments; 540nm/480nm for receptor internalization and FSH-mNG binding experiments.

## 5 Data availability

All the dataset and codes used are available in the following link: https://gitlab.inria.fr/cweckel/spatiotemporal_modeling2.

The scheme (Figs. 3-EV2) were done on MatchaIO.

## 6 Declaration of generative AI and AI-assisted technologies

During the preparation of this work the authors used Perplexity in order to improve readability. After using this tool, the authors reviewed and edited the content as needed, and they take full responsibility for the content of the published article.

## 7 Acknowledgements

This research was funded with the support of the Institut National de la Recherche pour l’Agriculture, l’Alimentation et l’Environnement (INRAE), INRAE Metaprogramme DIGIT-BIO (Digital biology to explore and predict living organisms in their environment), the Institut national de recherche en informatique et en automatique (INRIA), INRIA exploratory action (AEX) Compartimentage, the MAbImprove Labex (ANR-10-LABX-0053), the Région Centre Val de Loire ARD2020 Biomédicaments SELMAT grant, the Bill & Melinda Gates Foundation CONTRABODY grant, the ANR-22-CE14-0050 MOSDER grant and the PEPR Infertil-SaFe project code: ANR-24-PESF-0004.

J.G. was funded by a joint fellowship from MabImprove Labex and Bill & Melinda Gates foundation. C.W. was funded by a joint fellowship from INRAE and INRIA.

This work benefited from the IT infrastructure of the ISLANDe platform, particularly a computing cluster financed by the European Regional Development Fund n^°^ 159037.

---

**Figure EV 1.**
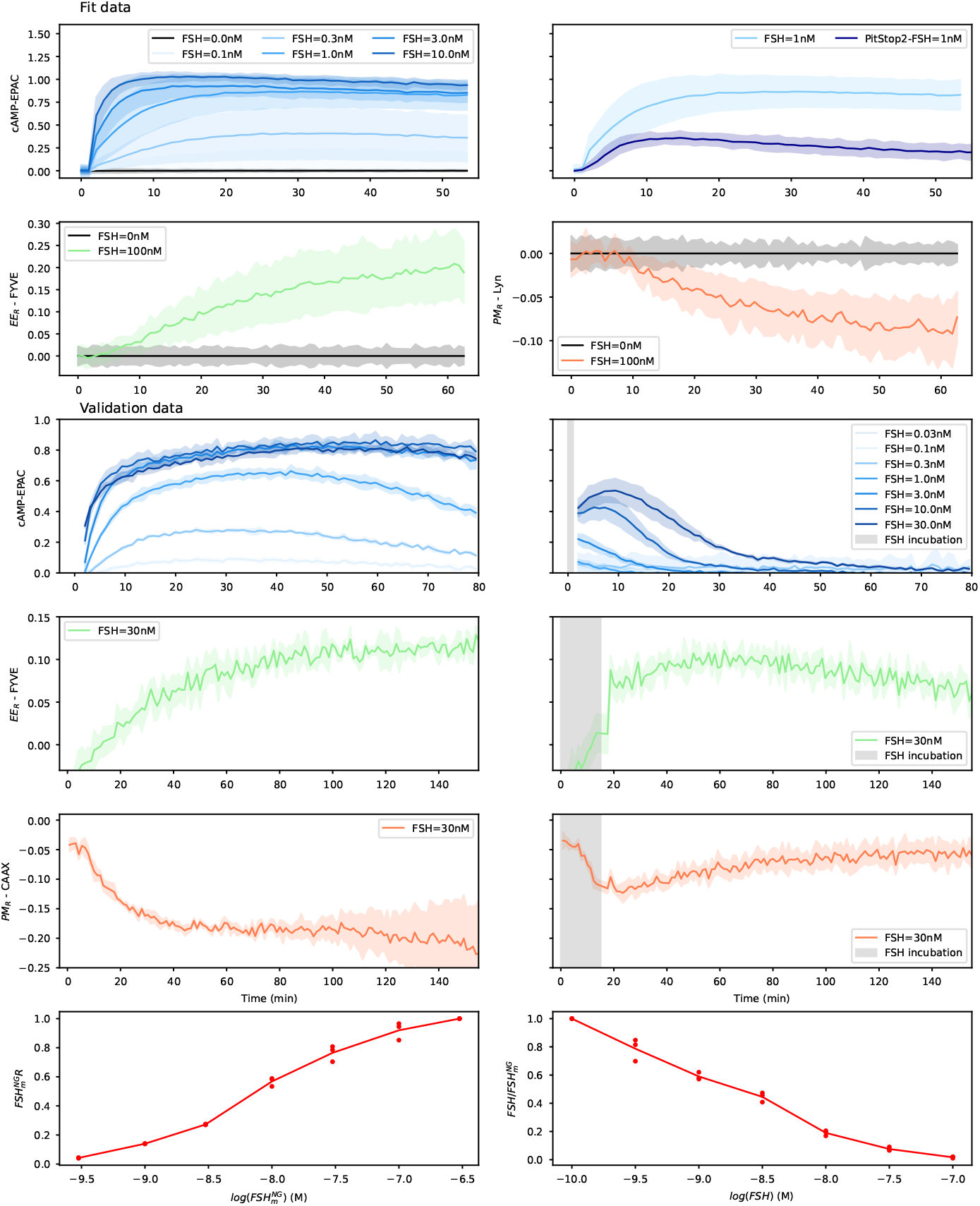
Data used for the fit and validation. For the fit, 10 replicates were used for each dataset. Four data types were included: cAMP levels measured with NLuc-EpacD602A-VV sensor (EPAC) at six ligand concentrations (0, 0.1, 0.3, 1, 3, 10 nM) and with CME inhibitor, PitStop2 for *FSH* = 1 nM, as well as plasma membrane (Lyn) and early endosomes (FYVE) localized receptors at two ligand concentrations (0 and 100 nM). For validation, continuous ligand stimulation is shown in the left column and pulse-chase in the right. cAMP (EPAC) is measured at seven doses (0, 0.01, 0.3, 1, 3, 10, 30 nM), along with receptor trafficking (CAAX and FYVE) at *L* = 30*nM* . Additional validation data include 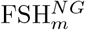 binding to FSHR at different concentrations and its competition with native FSH for receptor binding 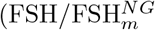.

**Figure EV 2.**
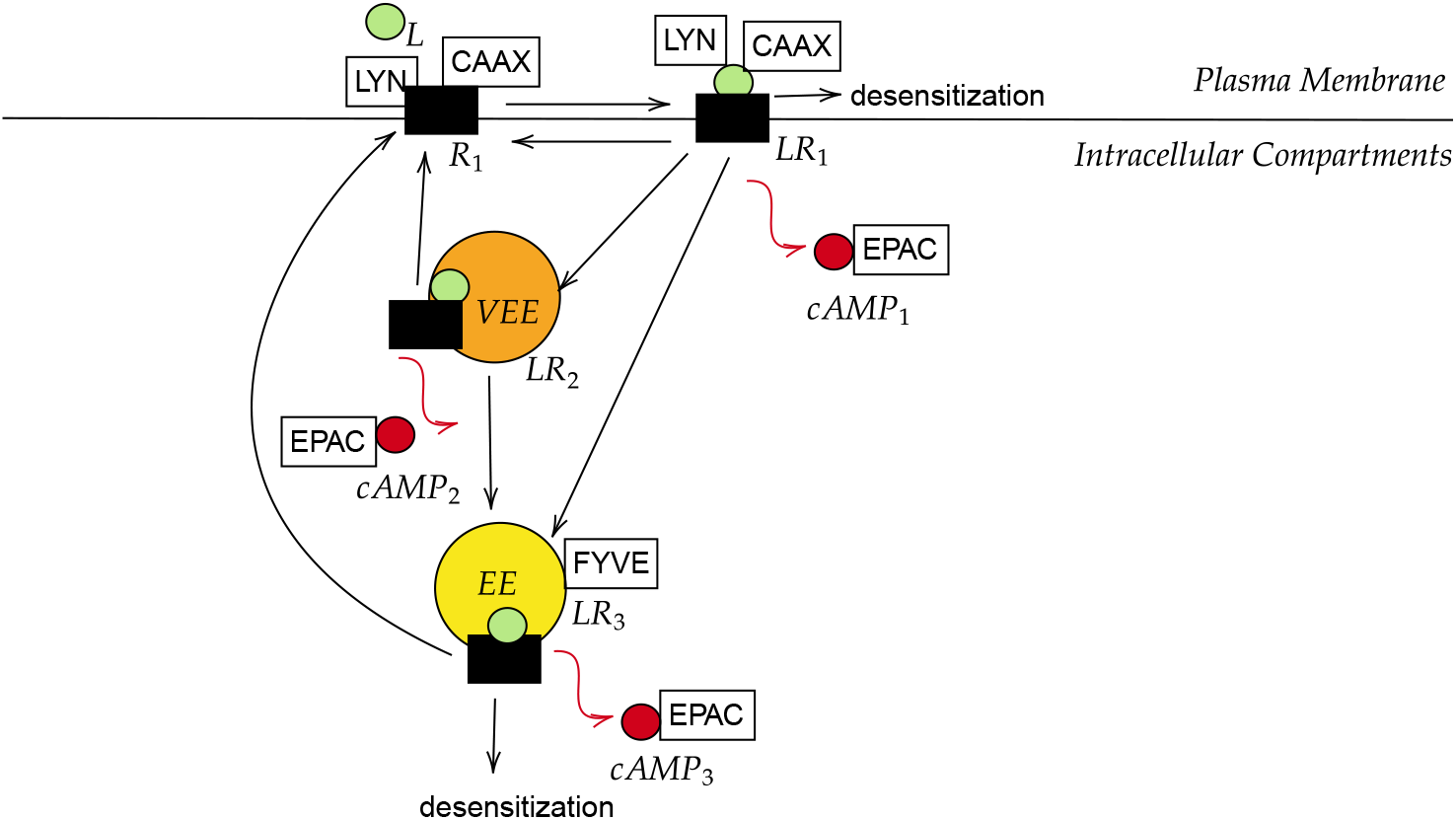
Biological Scheme. *R*_1_ represents the amount of receptors at the plasma membrane and *LR*_1_ (respectively *LR*_2_ and *LR*_3_) the amount of complex ligand-receptor at the plasma membrane (respectively in two types of intracellular compartments: very early endosomes and early endosomes). BRET sensing systems are represented in boxed texts: EPAC (cAMP measurement), CAAX and Lyn (receptor localized at plasma membrane) and FYVE (receptors localized at early endosomes).

**Figure EV 3.**
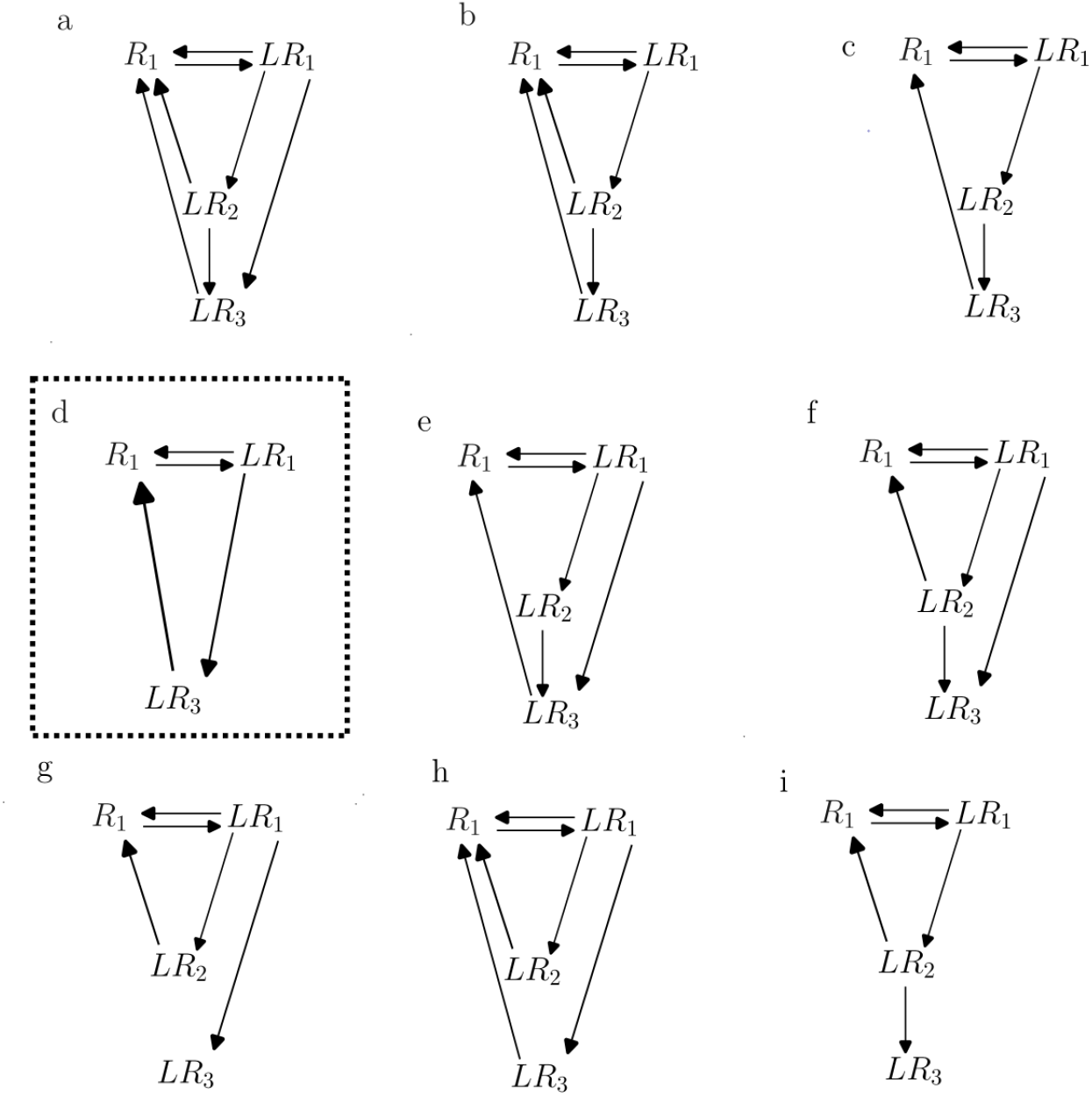
Trafficking models used for model selection. *R*_1_ represents the amount of receptors at the plasma membrane and *LR*_1_ (respectively *LR*_2_ and *LR*_3_) the amount of ligand-receptor complex at plasma membrane (respectively in two types of intracellular compartments). Model *d* corresponds to model with only two compartments (Q1), while all other models integrate three compartments. Then, different assumptions about trafficking are tested (a, b, c, e, f, g, h and i) (Q6): internalization of *LR*_1_, recycling of *LR*_2_ and *LR*_3_, trafficking between compartments (compartment 2 to compartment 3). All these models are embedded inside the complete three compartments model (Model 3). When no arrow is shown, the corresponding parameter value is set to zero.

**Figure EV 4.**
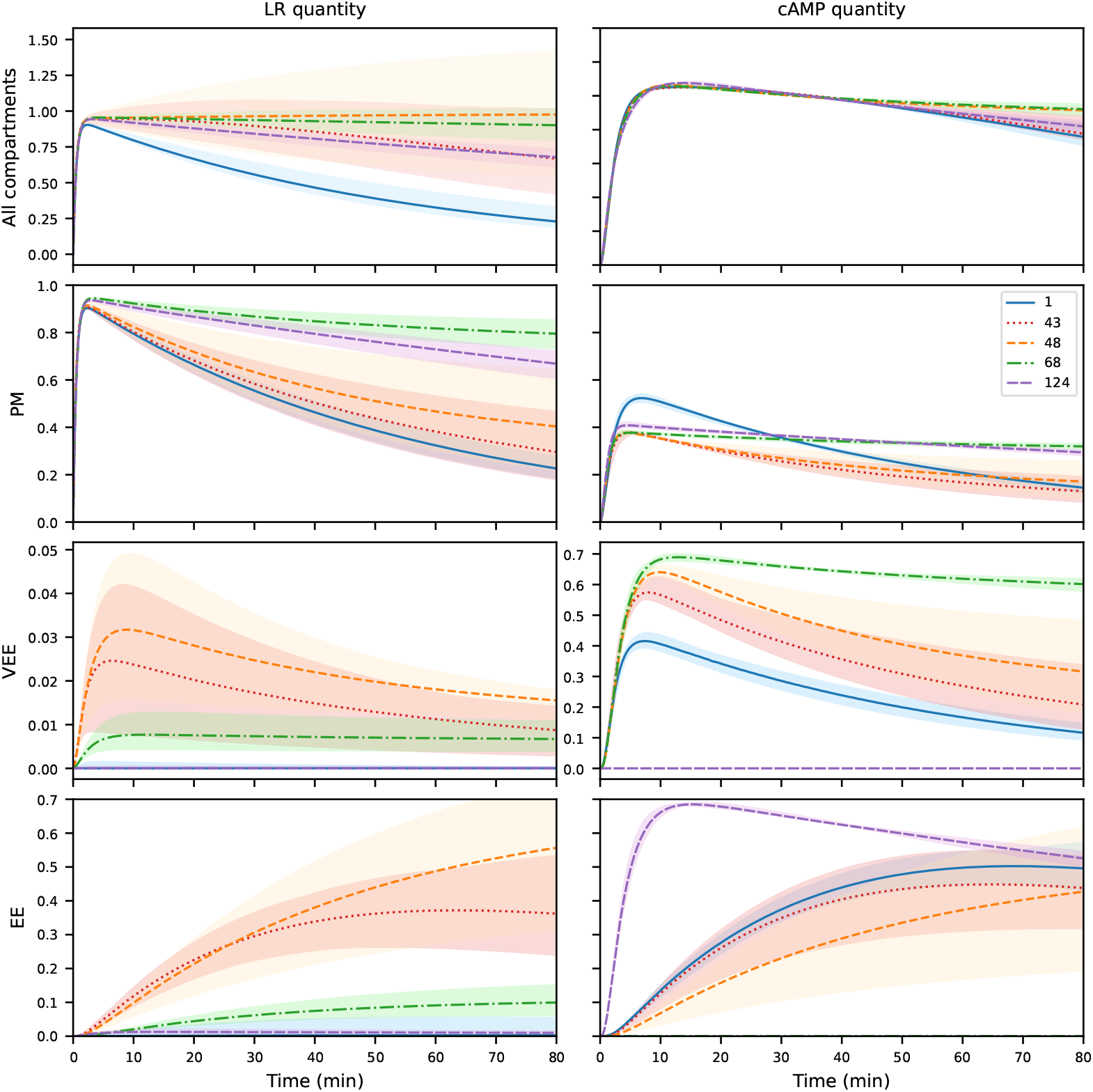
Prediction of cAMP levels and LR complex quantities in each compartment for four models. LR complex levels (first column) and cAMP quantities (second column) are predicted in each compartment for a single concentration of FSH (10 nM) for trafficking model *c*, using five different models (according to legend). The solid line (blue) represents the best model with the favorite cluster (3.1); the dotted line (red) corresponds to trafficking model *c* without desensitization at the plasma membrane but with desensitization from the compartment 3 (*LR*_3_) (3.43), the dashed line (orange) corresponds to trafficking model *c* without desensitization at the plasma membrane (3.48), the dash-dot line (green) represents trafficking model *c* with cAMP production restricted to the first two compartments (3.68), and the purple represents trafficking model *c* with cAMP production restricted to the two compartments *LR*_1_ and *LR*_3_ (3.124). Predictions are shown with 95% profile-based confidence intervals. The parameters used for these predictions correspond to the best estimates obtained in each parameter estimation problem (Appendix Table S4).

**Figure EV 5.**
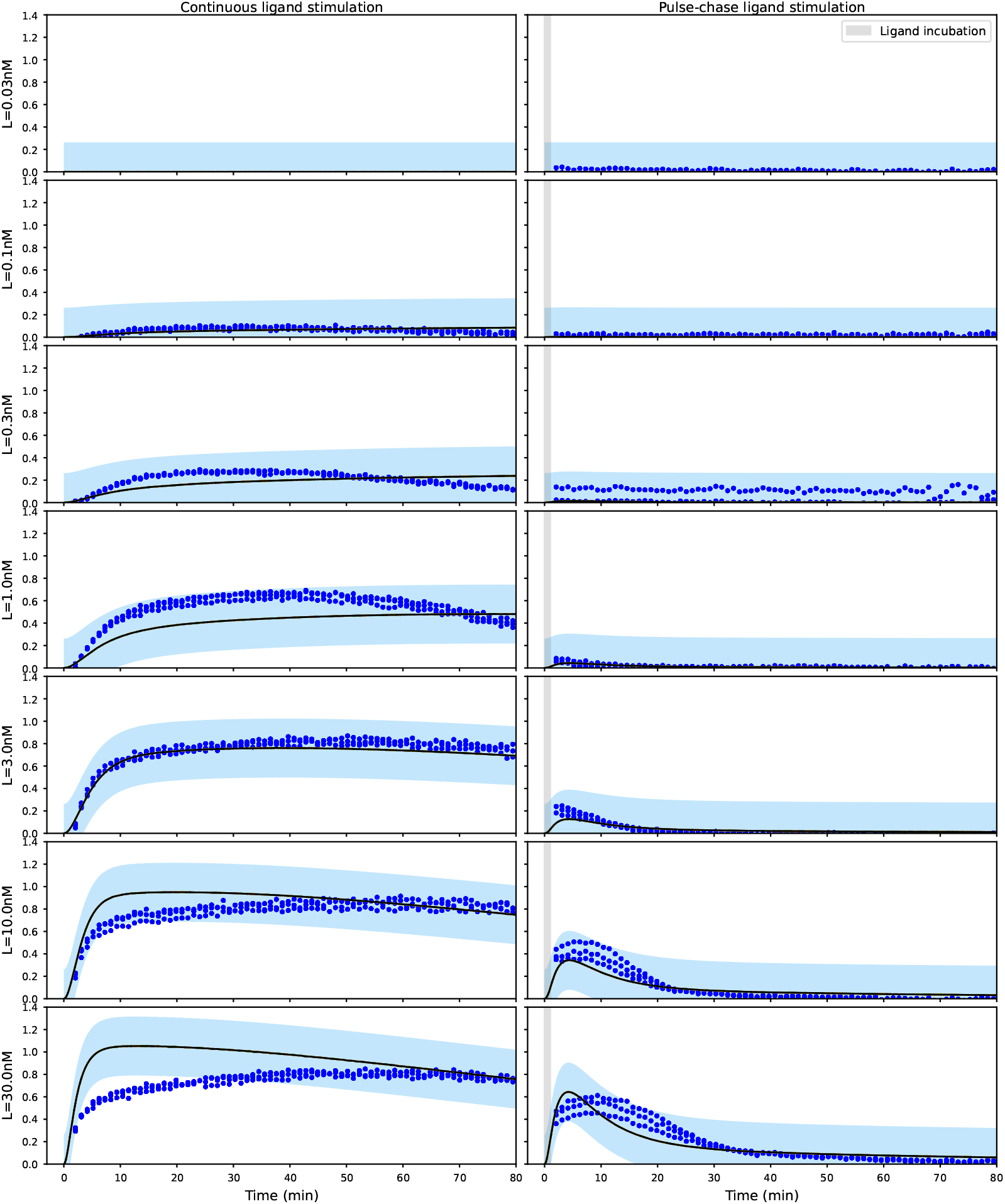
Prediction of cAMP quantity with validation data for the 13 best models. Continuous ligand stimulation condition is shown in the first column and pulse-chase ligand stimulation condition in the second column. cAMP quantity produced for 7 doses of ligand is showed. The parameters used for these predictions correspond to the best estimates obtained for the thirteen best models which were selected according to the AIC criterion (Appendix Table S 3) with their statistical error (shaded areas).

## 8 Supplementary Material

**Appendix Table S 1.**
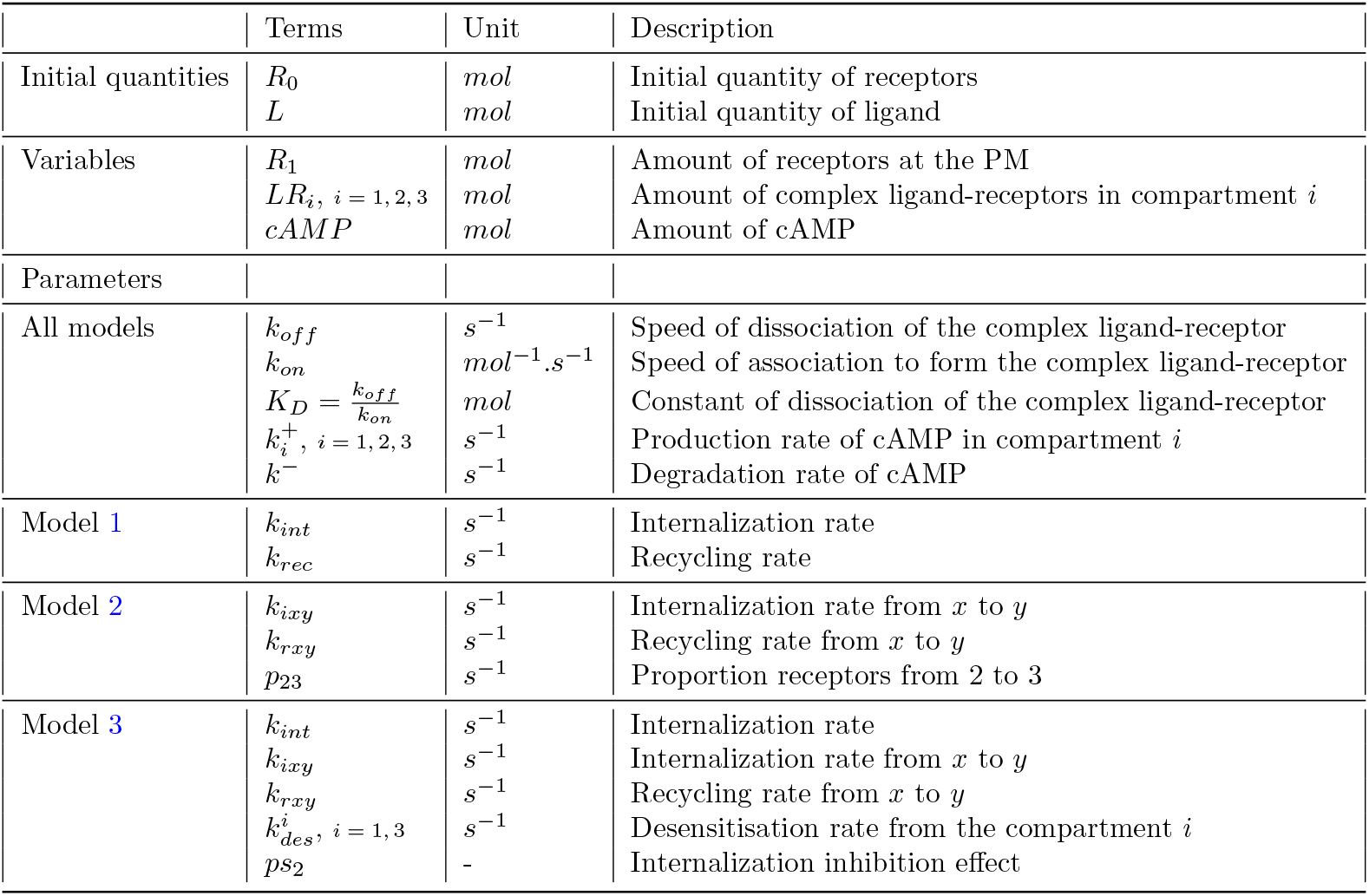
Table of the initial quantities, the variables and the parameters for models 1-2-3.

|  | Terms | Unit | Description |
| --- | --- | --- | --- |
| Initial quantities | $R_0$<br>$L$ | $mol$<br>$mol$ | Initial quantity of receptors<br>Initial quantity of ligand |
| Variables | $R_1$<br>$LR_i, i = 1, 2, 3$<br>$cAMP$ | $mol$<br>$mol$<br>$mol$ | Amount of receptors at the PM<br>Amount of complex ligand-receptors in compartment $i$<br>Amount of cAMP |
| Parameters |  |  |  |
| All models | $k_{off}$<br>$k_{on}$<br>$K_D = \frac{k_{off}}{k_{on}}$<br>$k_i^+, i = 1, 2, 3$<br>$k^-$ | $s^{-1}$<br>$mol^{-1}.s^{-1}$<br>$mol$<br>$s^{-1}$<br>$s^{-1}$ | Speed of dissociation of the complex ligand-receptor<br>Speed of association to form the complex ligand-receptor<br>Constant of dissociation of the complex ligand-receptor<br>Production rate of cAMP in compartment $i$<br>Degradation rate of cAMP |
| Model 1 | $k_{int}$<br>$k_{rec}$ | $s^{-1}$<br>$s^{-1}$ | Internalization rate<br>Recycling rate |
| Model 2 | $k_{ixy}$<br>$k_{rxy}$<br>$p_{23}$ | $s^{-1}$<br>$s^{-1}$<br>$s^{-1}$ | Internalization rate from $x$ to $y$<br>Recycling rate from $x$ to $y$<br>Proportion receptors from 2 to 3 |
| Model 3 | $k_{int}$<br>$k_{ixy}$<br>$k_{rxy}$<br>$k_{des}^i, i = 1, 3$<br>$ps_2$ | $s^{-1}$<br>$s^{-1}$<br>$s^{-1}$<br>$s^{-1}$<br>- | Internalization rate<br>Internalization rate from $x$ to $y$<br>Recycling rate from $x$ to $y$<br>Desensitisation rate from the compartment $i$<br>Internalization inhibition effect |

**Appendix Table S 2.** Result of model selection based on AIC criterion. 200 models were compared through several hypotheses (Subsect. 2.2.2): the number of compartments (Q1); whether the ligand-receptor complex is functionally active in two or three compartments (1 means active in the three compartments, 2 in *LR*_1_ and *LR*_2_ and 3 in *LR*_1_ and *LR*_3_) (Q2); if the receptor internalization inhibition drug is partially or fully efficient (Q3); if the receptor is desensitized at plasma membrane (Q4) or at early endosomes (Q5) and the different trafficking hypotheses (Q6). Crosses mean that the parameter is estimated, whereas when the box is empty, the corresponding parameter is fixed to 0.(Model 3)

| Model | H1 | H2 | H3 | H4 | H5 | H6 | AIC | $\Delta_{AIC}$ |
| --- | --- | --- | --- | --- | --- | --- | --- | --- |
| 1 | 3 | 1 |  | x |  | c | -16622.29 | 0.00 |
| 2 | 3 | 3 |  | x |  | e | -16622.22 | 0.07 |
| 3 | 3 | 1 |  | x |  | b | -16620.38 | 1.91 |
| 4 | 3 | 1 |  | x | x | i | -16620.38 | 1.91 |
| 5 | 3 | 1 |  | x |  | h | -16620.37 | 1.92 |
| 6 | 3 | 1 |  | x | x | g | -16620.36 | 1.92 |
| 7 | 3 | 1 |  | x | x | c | -16620.29 | 2.00 |
| 8 | 3 | 1 | x | x |  | c | -16620.29 | 2.00 |
| 9 | 3 | 1 |  | x |  | e | -16620.27 | 2.01 |
| 10 | 3 | 3 |  | x |  | a | -16620.22 | 2.07 |
| 11 | 3 | 3 |  | x | x | e | -16620.22 | 2.07 |
| 12 | 3 | 3 | x | x |  | e | -16620.22 | 2.07 |
| 13 | 3 | 3 |  | x | x | f | -16620.14 | 2.14 |
| 14 | 3 | 1 |  | x | x | b | -16618.38 | 3.91 |
| 15 | 3 | 1 | x | x |  | b | -16618.38 | 3.91 |
| 16 | 3 | 1 | x | x | x | i | -16618.38 | 3.91 |
| 17 | 3 | 1 | x | x |  | h | -16618.37 | 3.92 |
| 18 | 3 | 1 |  | x |  | a | -16618.37 | 3.92 |
| 19 | 3 | 1 |  | x | x | h | -16618.37 | 3.92 |
| 20 | 3 | 1 |  | x | x | f | -16618.36 | 3.92 |
| 21 | 3 | 1 | x | x | x | g | -16618.36 | 3.93 |
| 22 | 3 | 1 | x | x | x | c | -16618.29 | 4.00 |
| 23 | 3 | 1 |  | x | x | e | -16618.27 | 4.01 |
| 24 | 3 | 1 | x | x |  | e | -16618.27 | 4.01 |
| 25 | 3 | 3 |  | x | x | a | -16618.22 | 4.07 |
| 26 | 3 | 3 | x | x |  | a | -16618.22 | 4.07 |
| 27 | 3 | 3 | x | x | x | e | -16618.22 | 4.07 |
| 28 | 3 | 3 | x | x | x | f | -16618.14 | 4.14 |
| 29 | 3 | 1 | x | x | x | b | -16616.38 | 5.91 |
| 30 | 3 | 1 | x | x |  | a | -16616.37 | 5.92 |
| 31 | 3 | 1 | x | x | x | h | -16616.36 | 5.92 |
| 32 | 3 | 1 | x | x | x | f | -16616.36 | 5.92 |
| 33 | 3 | 1 |  | x | x | a | -16616.36 | 5.93 |
| 34 | 3 | 1 | x | x | x | e | -16616.27 | 6.02 |
| 35 | 3 | 3 | x | x | x | a | -16616.22 | 6.07 |
| 36 | 3 | 1 | x | x | x | a | -16614.36 | 7.93 |
| 37 | 3 | 1 |  | x |  | i | -16612.29 | 9.99 |
| 38 | 3 | 1 |  | x |  | g | -16612.28 | 10.01 |
| 39 | 3 | 1 | x | x |  | i | -16610.29 | 11.99 |
| 40 | 3 | 1 |  | x |  | f | -16610.28 | 12.01 |
| 41 | 3 | 1 | x | x |  | g | -16610.27 | 12.01 |
| 42 | 3 | 1 | x | x |  | f | -16608.27 | 14.01 |
| 43 | 3 | 1 |  |  | x | c | -16535.87 | 86.42 |
| 44 | 3 | 1 |  |  | x | i | -16533.88 | 88.41 |
| 45 | 3 | 1 |  |  | x | b | -16533.87 | 88.42 |
| 46 | 3 | 1 | x |  | x | c | -16533.86 | 88.42 |
| 47 | 3 | 1 |  |  | x | e | -16533.86 | 88.43 |
| 48 | 3 | 1 |  |  |  | c | -16532.15 | 90.14 |
| 49 | 3 | 1 | x |  | x | i | -16531.88 | 90.41 |
| 50 | 3 | 1 |  |  | x | f | -16531.88 | 90.41 |
| 51 | 3 | 1 | x |  | x | b | -16531.86 | 90.42 |
| 52 | 3 | 1 |  |  | x | a | -16531.86 | 90.43 |
| 53 | 3 | 1 | x |  | x | e | -16531.86 | 90.43 |
| 54 | 3 | 1 |  |  |  | b | -16530.15 | 92.14 |
| 55 | 3 | 1 | x |  |  | c | -16530.15 | 92.14 |
| 56 | 3 | 1 |  |  |  | e | -16530.15 | 92.14 |
| 57 | 3 | 1 | x |  | x | f | -16529.87 | 92.41 |
| 58 | 3 | 1 | x |  | x | a | -16529.86 | 92.43 |
| 59 | 3 | 1 | x |  |  | b | -16528.15 | 94.14 |
| 60 | 3 | 1 |  |  |  | a | -16528.15 | 94.14 |
| 61 | 3 | 1 | x |  |  | e | -16528.14 | 94.14 |
| 62 | 3 | 1 |  |  | x | g | -16527.05 | 95.23 |
| 63 | 3 | 1 | x |  |  | a | -16526.14 | 96.14 |
| 64 | 3 | 1 |  |  | x | h | -16525.35 | 96.93 |
| 65 | 3 | 1 | x |  | x | g | -16525.05 | 97.24 |
| 66 | 3 | 1 |  |  |  | i | -16525.01 | 97.28 |
| 67 | 3 | 2 |  |  |  | c | -16523.89 | 98.40 |
| 68 | 3 | 2 |  | x |  | c | -16523.68 | 98.61 |
| 69 | 3 | 1 | x |  | x | h | -16523.35 | 98.94 |
| 70 | 3 | 1 |  |  |  | f | -16523.01 | 99.28 |
| 71 | 3 | 1 | x |  |  | i | -16523.01 | 99.28 |
| 72 | 3 | 2 |  |  |  | b | -16521.89 | 100.40 |
| 73 | 3 | 2 | x |  |  | c | -16521.89 | 100.40 |
| 74 | 3 | 2 |  |  | x | c | -16521.88 | 100.41 |
| 75 | 3 | 2 |  |  |  | e | -16521.87 | 100.41 |
| 76 | 3 | 2 |  | x |  | b | -16521.68 | 100.61 |
| 77 | 3 | 2 | x | x |  | c | -16521.68 | 100.61 |
| 78 | 3 | 2 |  | x | x | c | -16521.67 | 100.62 |
| 79 | 3 | 2 |  | x |  | e | -16521.66 | 100.62 |
| 80 | 3 | 1 | x |  |  | f | -16521.01 | 101.28 |
| 81 | 3 | 2 | x |  |  | b | -16519.89 | 102.40 |
| 82 | 3 | 2 |  |  | x | b | -16519.88 | 102.41 |
| 83 | 3 | 2 | x |  | x | c | -16519.88 | 102.41 |
| 84 | 3 | 2 |  |  |  | a | -16519.87 | 102.41 |
| 85 | 3 | 2 | x |  |  | e | -16519.87 | 102.41 |
| 86 | 3 | 2 |  |  | x | e | -16519.87 | 102.42 |
| 87 | 3 | 2 | x | x |  | b | -16519.68 | 102.61 |
| 88 | 3 | 2 |  | x | x | b | -16519.67 | 102.62 |
| 89 | 3 | 2 | x | x | x | c | -16519.67 | 102.62 |
| 90 | 3 | 2 |  | x |  | a | -16519.66 | 102.62 |
| 91 | 3 | 2 | x | x |  | e | -16519.66 | 102.62 |
| 92 | 3 | 2 |  | x | x | e | -16519.66 | 102.63 |
| 93 | 3 | 2 | x |  | x | b | -16517.88 | 104.41 |
| 94 | 3 | 2 | x |  |  | a | -16517.87 | 104.41 |
| 95 | 3 | 2 |  |  | x | a | -16517.87 | 104.42 |
| 96 | 3 | 2 | x |  | x | e | -16517.87 | 104.42 |
| 97 | 3 | 2 | x | x | x | b | -16517.67 | 104.62 |
| 98 | 3 | 2 | x | x |  | a | -16517.66 | 104.62 |
| 99 | 3 | 2 |  | x | x | a | -16517.66 | 104.63 |
| 100 | 3 | 2 | x | x | x | e | -16517.66 | 104.63 |
| 101 | 3 | 1 |  |  |  | h | -16517.56 | 104.72 |
| 102 | 3 | 1 |  |  |  | g | -16516.64 | 105.65 |
| 103 | 3 | 2 | x |  | x | a | -16515.87 | 106.42 |
| 104 | 3 | 2 | x | x | x | a | -16515.66 | 106.63 |
| 105 | 3 | 1 | x |  |  | h | -16515.56 | 106.72 |
| 106 | 3 | 1 | x |  |  | g | -16514.63 | 107.65 |
| 107 | 3 | 2 |  | x |  | h | -16511.10 | 111.19 |
| 108 | 3 | 2 | x | x |  | h | -16509.10 | 113.19 |
| 109 | 3 | 2 |  | x | x | h | -16509.09 | 113.19 |
| 110 | 3 | 2 |  |  |  | h | -16508.98 | 113.31 |
| 111 | 3 | 2 | x | x | x | h | -16507.09 | 115.19 |
| 112 | 3 | 2 | x |  |  | h | -16506.98 | 115.31 |
| 113 | 3 | 2 |  |  | x | h | -16506.97 | 115.31 |
| 114 | 3 | 2 | x |  | x | h | -16504.97 | 117.31 |
| 115 | 3 | 3 |  | x |  | h | -16460.68 | 161.61 |
| 116 | 3 | 3 |  | x | x | h | -16458.68 | 163.61 |
| 117 | 3 | 3 | x | x |  | h | -16458.68 | 163.61 |
| 118 | 3 | 3 | x | x | x | h | -16456.68 | 165.61 |
| 119 | 2 | 3 |  | x |  | d | -16456.22 | 166.07 |
| 120 | 3 | 3 |  | x | x | g | -16455.99 | 166.30 |
| 121 | 3 | 3 |  |  | x | h | -16455.47 | 166.82 |
| 122 | 2 | 3 | x | x |  | d | -16454.22 | 168.07 |
| 123 | 2 | 3 |  | x | x | d | -16454.22 | 168.07 |
| 124 | 3 | 3 |  | x |  | c | -16454.08 | 168.21 |
| 125 | 3 | 3 | x | x | x | g | -16453.99 | 168.30 |
| 126 | 3 | 3 | x |  | x | h | -16453.47 | 168.82 |
| 127 | 3 | 3 |  |  | x | e | -16452.30 | 169.99 |
| 128 | 2 | 3 | x | x | x | d | -16452.22 | 170.07 |
| 129 | 3 | 3 |  | x |  | b | -16452.08 | 170.21 |
| 130 | 3 | 3 | x | x |  | c | -16452.08 | 170.21 |
| 131 | 3 | 3 |  | x | x | c | -16452.08 | 170.21 |
| 132 | 3 | 3 |  |  | x | a | -16450.30 | 171.99 |
| 133 | 3 | 3 | x |  | x | e | -16450.30 | 171.99 |
| 134 | 3 | 3 | x | x |  | b | -16450.08 | 172.21 |
| 135 | 3 | 3 |  | x | x | b | -16450.08 | 172.21 |
| 136 | 3 | 3 | x | x | x | c | -16450.08 | 172.21 |
| 137 | 3 | 3 | x |  | x | a | -16448.30 | 173.99 |
| 138 | 3 | 3 | x | x | x | b | -16448.08 | 174.21 |
| 139 | 3 | 2 |  | x |  | g | -16440.96 | 181.33 |
| 140 | 3 | 3 |  |  | x | g | -16440.81 | 181.48 |
| 141 | 3 | 2 |  | x |  | i | -16439.86 | 182.43 |
| 142 | 3 | 2 |  | x | x | g | -16438.96 | 183.33 |
| 143 | 3 | 2 | x | x |  | g | -16438.96 | 183.33 |
| 144 | 3 | 3 | x |  | x | g | -16438.81 | 183.48 |
| 145 | 3 | 3 |  |  | x | f | -16437.95 | 184.34 |
| 146 | 3 | 2 |  | x | x | i | -16437.86 | 184.43 |
| 147 | 3 | 2 | x | x |  | i | -16437.86 | 184.43 |
| 148 | 3 | 2 |  | x |  | f | -16437.79 | 184.49 |
| 149 | 3 | 2 | x | x | x | g | -16436.96 | 185.33 |
| 150 | 3 | 3 |  | x | x | i | -16436.46 | 185.83 |
| 151 | 3 | 3 | x |  | x | f | -16435.95 | 186.34 |
| 152 | 3 | 2 | x | x | x | i | -16435.86 | 186.43 |
| 153 | 3 | 2 |  | x | x | f | -16435.79 | 186.49 |
| 154 | 3 | 2 | x | x |  | f | -16435.79 | 186.49 |
| 155 | 3 | 3 | x | x | x | i | -16434.57 | 187.72 |
| 156 | 3 | 2 | x | x | x | f | -16433.79 | 188.49 |
| 157 | 2 | 3 |  |  | x | d | -16419.05 | 203.23 |
| 158 | 2 | 3 | x |  | x | d | -16417.05 | 205.23 |
| 159 | 3 | 3 |  |  | x | c | -16417.01 | 205.28 |
| 160 | 3 | 3 |  |  | x | b | -16415.01 | 207.28 |
| 161 | 3 | 3 | x |  | x | c | -16415.01 | 207.28 |
| 162 | 3 | 3 | x |  | x | b | -16413.01 | 209.28 |
| 163 | 3 | 2 |  |  |  | g | -16404.06 | 218.23 |
| 164 | 3 | 2 |  |  | x | g | -16402.06 | 220.23 |
| 165 | 3 | 2 | x |  |  | g | -16402.06 | 220.23 |
| 166 | 3 | 2 |  |  |  | f | -16401.62 | 220.67 |
| 167 | 3 | 2 | x |  | x | g | -16400.06 | 222.23 |
| 168 | 3 | 2 |  |  | x | f | -16399.62 | 222.67 |
| 169 | 3 | 2 | x |  |  | f | -16399.62 | 222.67 |
| 170 | 3 | 2 | x |  | x | f | -16397.62 | 224.67 |
| 171 | 3 | 3 |  |  | x | i | -16396.79 | 225.50 |
| 172 | 3 | 3 | x |  | x | i | -16394.79 | 227.50 |
| 173 | 3 | 2 | x |  |  | i | -16389.08 | 233.21 |
| 174 | 3 | 2 | x |  | x | i | -16387.08 | 235.21 |
| 175 | 3 | 2 |  |  |  | i | -16380.23 | 242.06 |
| 176 | 3 | 2 |  |  | x | i | -16378.23 | 244.06 |
| 177 | 3 | 3 |  | x |  | g | -16157.25 | 465.04 |
| 178 | 3 | 3 |  | x |  | i | -16156.97 | 465.31 |
| 179 | 3 | 3 |  | x |  | f | -16155.25 | 467.04 |
| 180 | 3 | 3 | x | x |  | g | -16155.24 | 467.05 |
| 181 | 3 | 3 | x | x |  | i | -16154.97 | 467.32 |
| 182 | 3 | 3 | x | x |  | f | -16153.24 | 469.05 |
| 183 | 3 | 3 |  |  |  | e | -15673.45 | 948.84 |
| 184 | 3 | 3 |  |  |  | a | -15671.44 | 950.85 |
| 185 | 3 | 3 | x |  |  | e | -15671.44 | 950.85 |
| 186 | 3 | 3 | x |  |  | a | -15669.43 | 952.86 |
| 187 | 3 | 3 |  |  |  | h | -15563.76 | 1058.53 |
| 188 | 3 | 3 | x |  |  | h | -15561.75 | 1060.53 |
| 189 | 2 | 3 |  |  |  | d | -15064.93 | 1557.35 |
| 190 | 3 | 3 |  |  |  | g | -15063.02 | 1559.27 |
| 191 | 3 | 3 |  |  |  | i | -15062.99 | 1559.30 |
| 192 | 2 | 3 | x |  |  | d | -15062.91 | 1559.38 |
| 193 | 3 | 3 |  |  |  | c | -15062.88 | 1559.41 |
| 194 | 3 | 3 |  |  |  | f | -15061.04 | 1561.25 |
| 195 | 3 | 3 | x |  |  | g | -15060.99 | 1561.29 |
| 196 | 3 | 3 | x |  |  | i | -15060.96 | 1561.33 |
| 197 | 3 | 3 |  |  |  | b | -15060.88 | 1561.41 |
| 198 | 3 | 3 | x |  |  | c | -15060.85 | 1561.44 |
| 199 | 3 | 3 | x |  |  | f | -15059.01 | 1563.28 |
| 200 | 3 | 3 | x |  |  | b | -15058.85 | 1563.44 |

**Appendix Table S 3.**
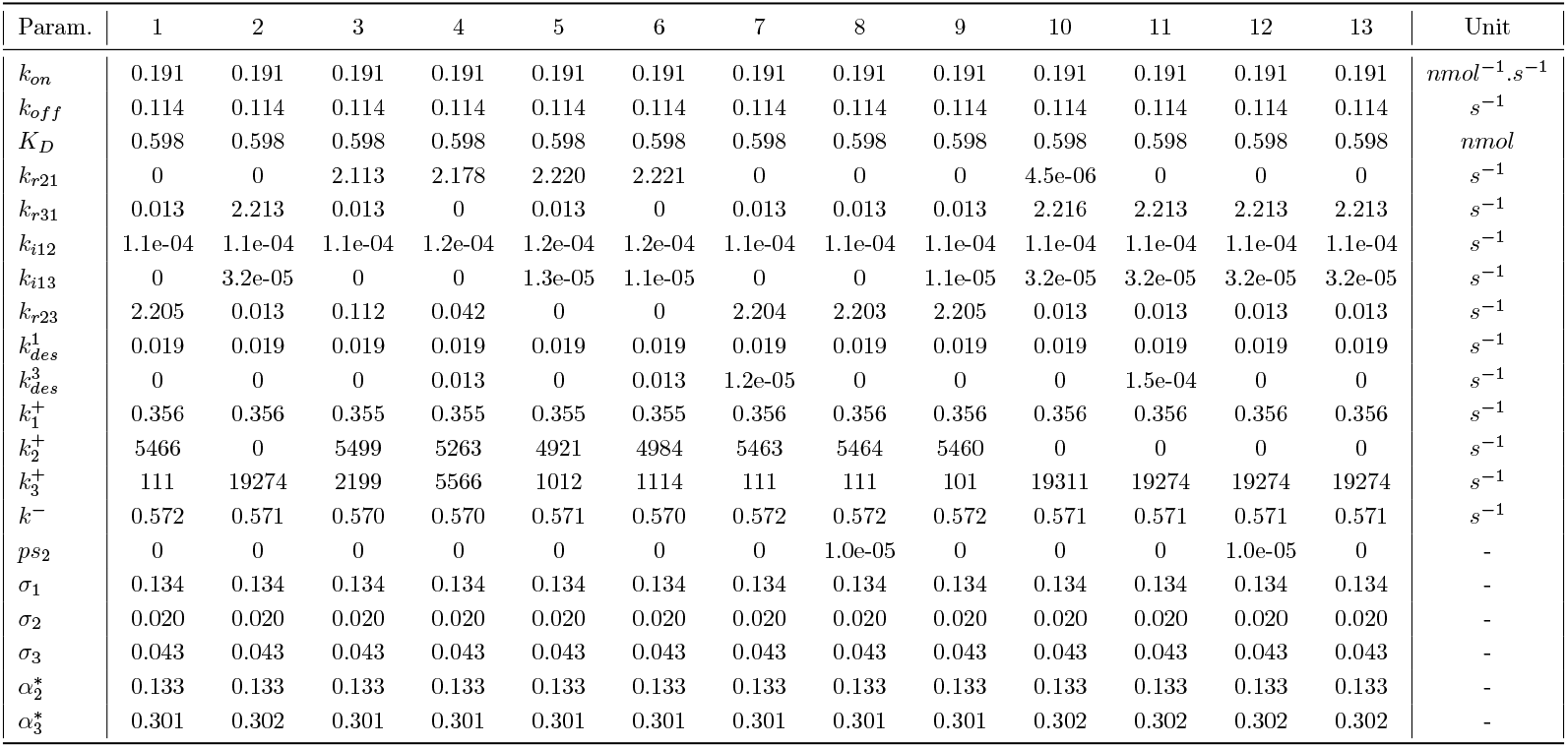
Table of the value of the best estimated parameters for the thirteen best models.

**Appendix Table S 4.**
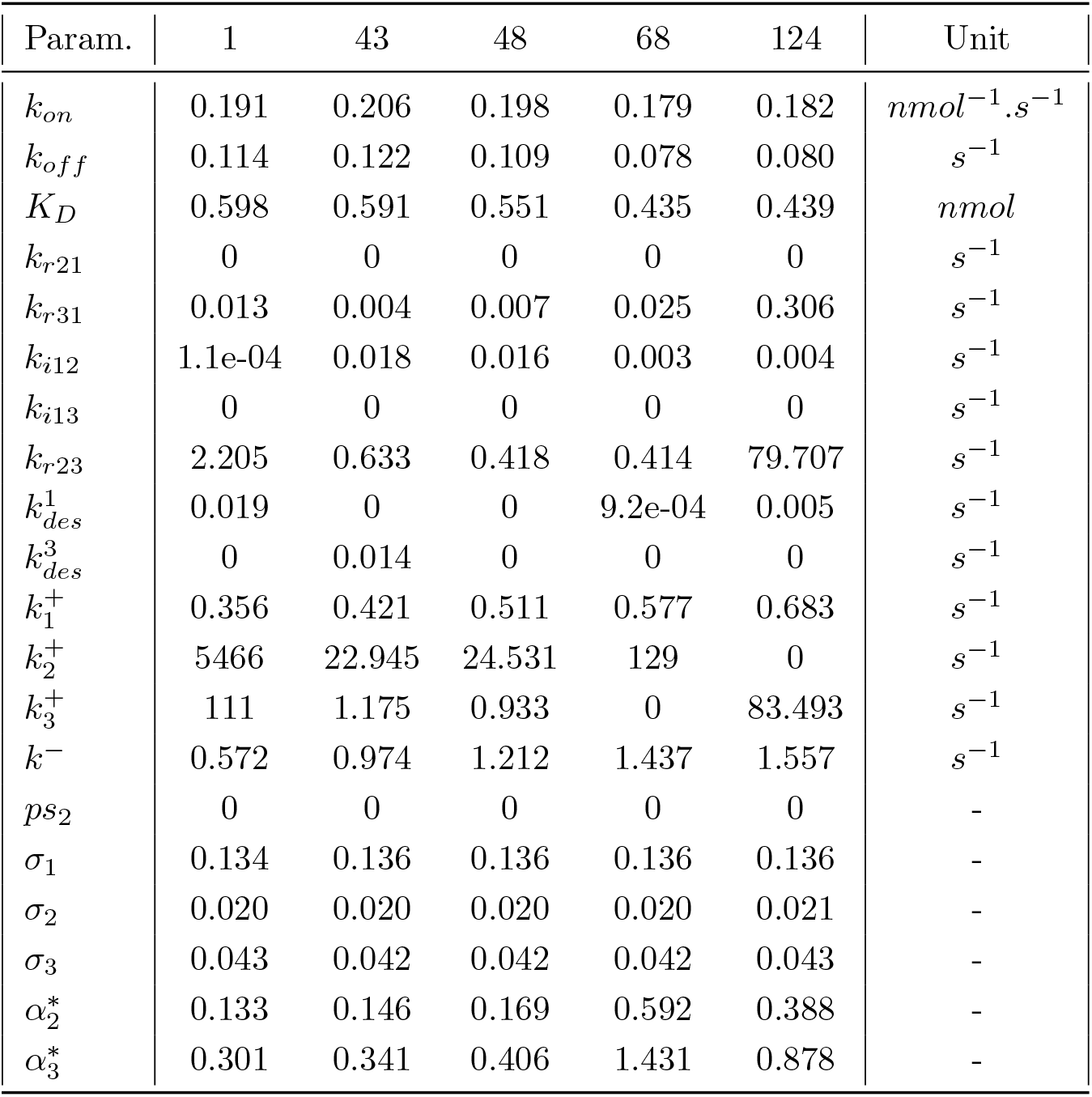
Table of the value of best parameters for five different models.

**Appendix Figure S 1.**
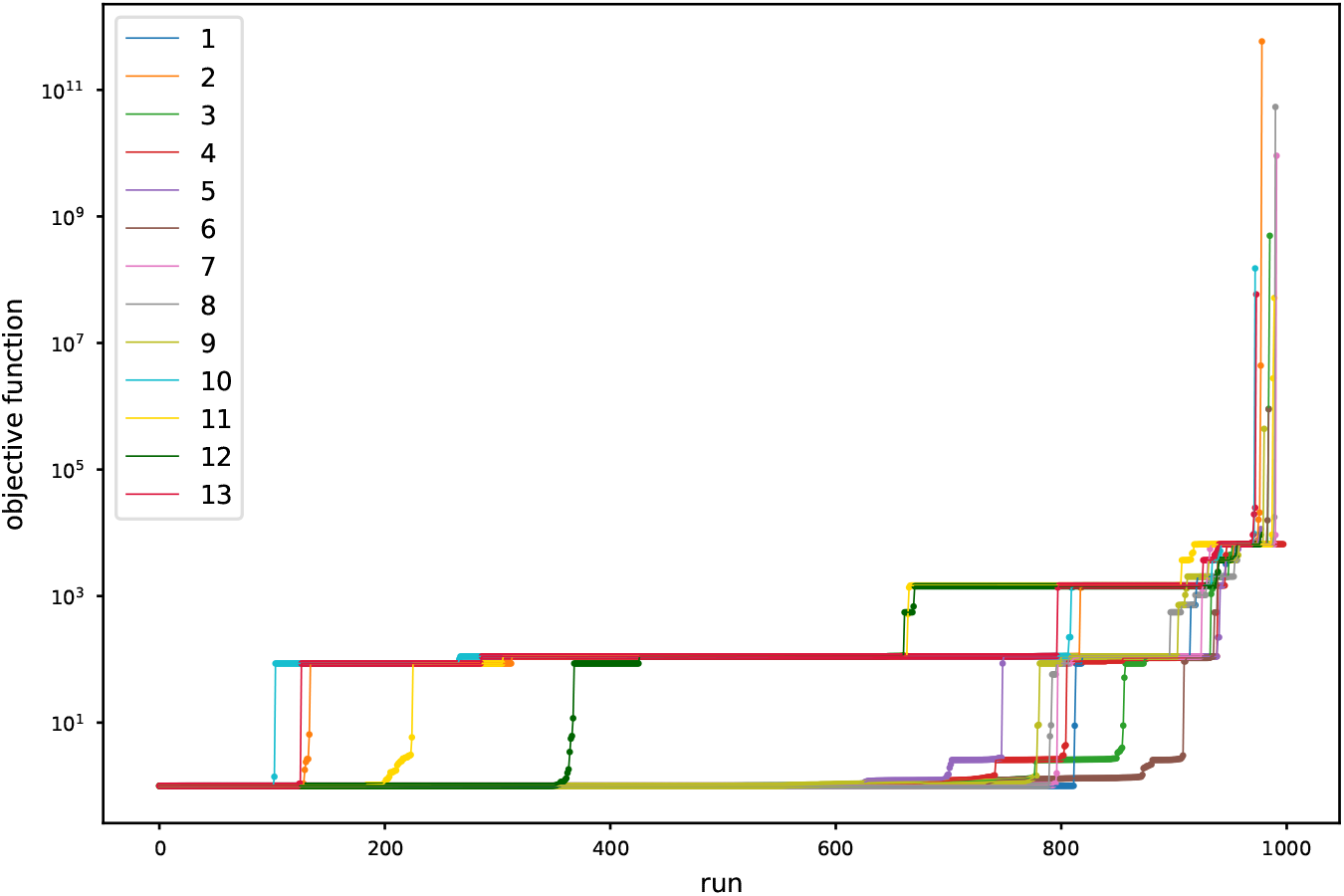
Convergence algorithm for the 13 models for the 1000 runs.

**Appendix Figure S 2.**
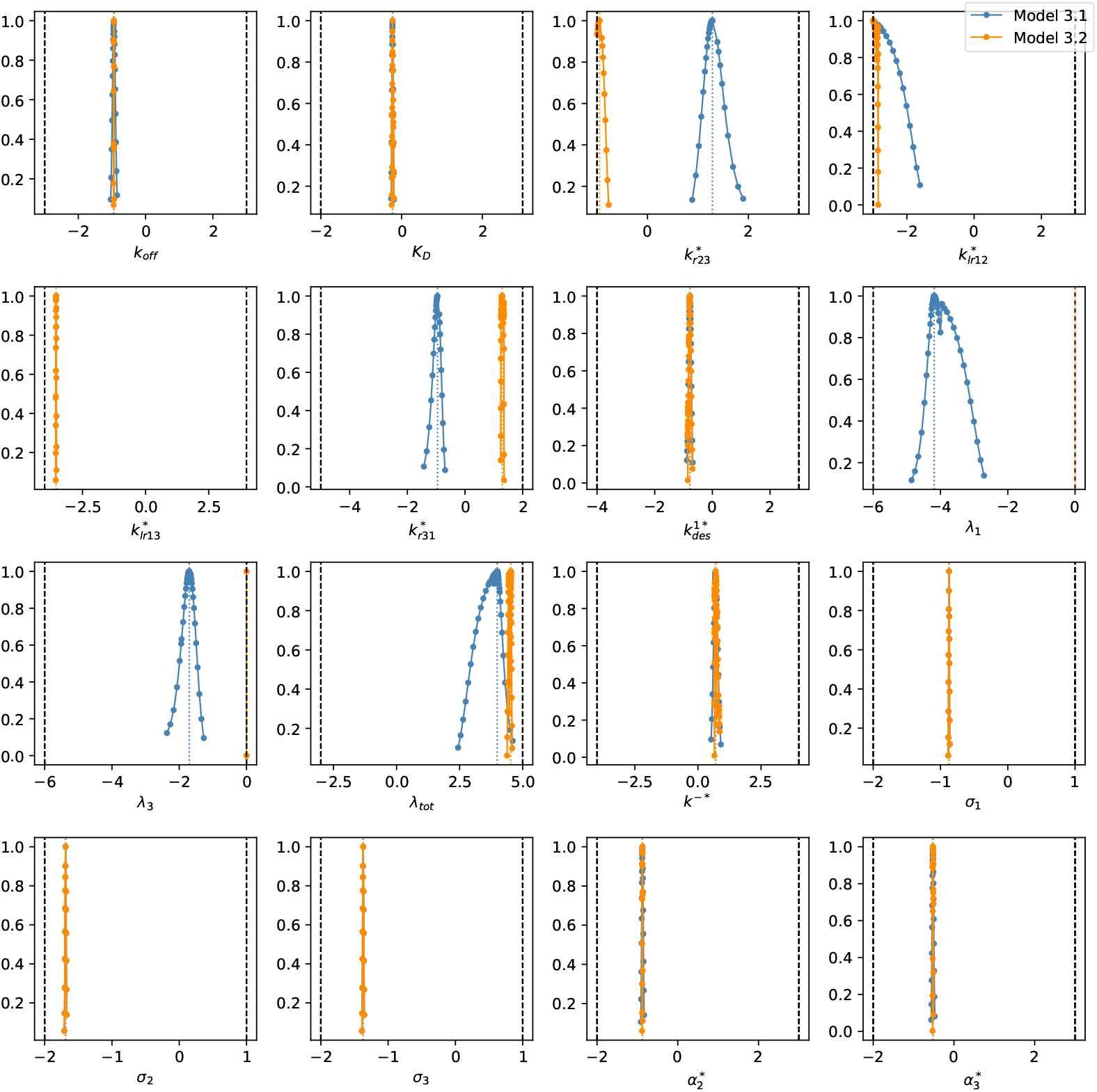
Profile likelihood for all parameters for models 3.1 and 3.2.

## Footnotes

1 Models 8 and 12 are the only two models that estimates the parameter *ps*_2_, but it does so with a very low value.

